# The Hemodynamic Response Function Varies Across Anatomical Location and Pathology in the Epileptic Brain

**DOI:** 10.64898/2025.12.16.694757

**Authors:** Zhengchen Cai, Nicolás von Ellenrieder, Thaera Arafat, Hui Ming Kho, Gang Chen, Andreas Koupparis, Chifaou Abdallah, Roy Dudley, Dang Khoa Nguyen, Jeffery Hall, Francois Dubeau, Jean Gotman, Boris Bernhardt

## Abstract

The hemodynamic response function (HRF) links neuronal activity to functional magnetic resonance imaging (fMRI) signals. While most fMRI studies use a “canonical” HRF, increasing evidence from studies of healthy subjects suggests that the HRF depends on anatomical location and disease states. Here, we investigate how HRF variability relates to anatomical location and pathology in the epileptic brain, using a large simultaneous electroencephalogram and fMRI dataset. Applying HRF deconvolution and temporal decomposition, we built the first whole-brain HRF library specific to epilepsy, identifying four distinct shape groups. We mapped HRF features across parcellations of two atlases using novel Bayesian hierarchical models. In non-epileptogenic regions, HRF shape and spatial distributions align with findings from healthy subjects. Within pathological regions, they vary significantly according to pathology. Our results indicate that HRF variability is associated with pathology, in addition to its dependence on anatomical location, motivating region- and pathology-based HRF modulation in epilepsy studies.

## Introduction

Epilepsy affects around 50 million people worldwide, but roughly one third of patients have seizures uncontrolled by medication. For many of these patients, surgery can be an effective treatment^1,2^. Successful surgery heavily relies on accurate localization of the epileptogenic zone. Simultaneous electroencephalography (EEG) and functional magnetic resonance imaging (fMRI) is a unique, noninvasive, and individualized multimodal technique that has proven effective for presurgical evaluation^3–9^. By detecting blood oxygen level dependent (BOLD) signal changes of fMRI associated with interictal epileptic discharges (IEDs, *e.g.,* spikes) identified from scalp EEG, EEG-fMRI enables direct mapping of patient specific epileptogenic regions and their propagation within the epileptic network^10–12^. The BOLD response to IEDs is mediated by the hemodynamic response function (HRF), which characterizes the temporal dynamics of neurovascular coupling as a nonlinear combination of cerebral blood flow and volume, and deoxygenated hemoglobin concentration changes^13,14^. FMRI analysis typically uses a “canonical” HRF derived from healthy brains, assuming a uniform shape across all brain regions and individuals. This can reduce both sensitivity and specificity in detecting BOLD responses, particularly when applied to estimate IED related responses in epilepsy studies.

Accumulating evidence shows that HRF varies temporally in peak amplitude, time and duration^15^, spatially across brain regions^16–18^, demographically with age^19^ and sex^20^, and experimentally across tasks^21^. Investigating this variability is crucial for accurate fMRI analysis, especially in epilepsy studies. Several approaches have been developed to incorporate HRF variability. While the approach using temporal derivatives and dispersion^22^ is more constrained (*e.g.,* peak time and width only vary ∼1s), more flexible data driven deconvolution methods^23–26^ suffer from overfitting. Testing multiple predefined HRFs and selecting the one providing the strongest statistical response offers a practical balance^27,28^, but they are typically defined by mathematical functions (*e.g.,* gamma function) rather than data informed shapes. Recently, Prince et al. (2022) further advanced this strategy by using empirically derived HRF shapes from the visual cortex of healthy subjects, demonstrated improved task trial level response estimation, enhanced between participant representational similarity across datasets and improved natural scene image decoding^29^. These findings raise interesting fundamental questions about HRF variability in epilepsy: specifically, how many representative HRFs can be identified in the epileptic brain, what are their temporal features, and how they differ from those in healthy subjects. HRF features also vary spatially across brain regions in healthy individuals^16–18^, but how this spatial variability manifests in the epileptic brain remains unknown. HRF variability has also been linked to neurological disorders. For instance, individuals with subjective cognitive decline showed altered HRF peak amplitude and latency compared to healthy controls^30^. In epilepsy, delayed HRF peaks have been reported in children under two years of age^31^, while adults with epilepsy exhibited wider, delayed HRFs during auditory motor tasks relative to the canonical HRF^32^. No significant differences in HRF peak amplitude or latency were identified between patients with focal cortical dysplasia and those with hippocampal sclerosis^33^. Yet, no study to date has systematically compared HRF features across different anatomical locations in the epileptic brain, especially with different pathologies.

Here, we comprehensively characterized the temporal and spatial features of HRF in the epileptic brain. Our investigation was based on a large and well-defined clinical dataset comprising 132 IED related fMRI response maps of 84 scans (*i.e.,* including 79 patients with multiple IED types) collected in pharmaco-resistant epilepsy patients who underwent presurgical EEG-fMRI. Notably, all included patients also underwent surgical resection, and the majority had a histopathological diagnosis subsequent to EEG-fMRI. Our work featured three primary studies: First, we built a data-driven HRF library specific to the epileptic brain using regularized HRF deconvolution^18^ and temporal decomposition^34^ methods, identifying representative HRF shapes that varied in amplitude and timing, which were further grouped into four distinct shape groups. Second, we mapped HRF features throughout the epileptic brain and investigated their associations with anatomical location using data from regions outside the cavities resected during subsequent surgeries. Third, we evaluated whether HRF features are associated with pathology using data from regions that were subsequently resected and histologically analyzed. To obtain more accurate, reliable and generalizable estimates of associations, we developed a novel hierarchical Bayesian model that performs partial pooling of HRF features following the brain’s anatomical hierarchy (parcel-lobe-brain), enabling estimated effects to be informed by both local and global homogeneity. We further demonstrated these associations through model comparison and machine learning based pathology classification using HRF features alone, and confirmed robustness of results across two different parcellation atlases. Altogether, our study provides the first HRF library covering the entire epileptic brain and reveals systematic HRF variation associated with pathology in addition to its dependence on anatomical location.

## Results

Here, we report the results following our three main studies: the newly generated HRF library; the demonstration of a clear association between HRF characteristics and anatomical location; and the demonstration of an association between HRF characteristics and pathology. These results are further supported by demonstrating robustness to the choice of parcellation atlas, and the effects of other factors such as age, gender, IED type, and HRF sign on HRF features. Model fittings under the Bayesian framework converged well, and the effective posterior sample size of every parameter was sufficient for accurate statistical summary based on recommended criteria^35,36^. We present each effect intuitively using the entire estimated distribution and summarize it numerically with the median and the 95% highest density interval (HDI_95%_) to characterize uncertainty.

### HRF deconvolution analysis of IED-related fMRI response

Our deconvolution analysis identified 132 IED-related fMRI response maps from 84 scans that showed 154,122 statistically significant voxels, using a two-sided p-value threshold of 0.001 for both the *F*-statistic of general linear model (GLM) model and the maximum *t*-value across all 17 (0 to 25.5s, 1.5s increment) deconvolution basis functions. The number of statistically significant voxels varied across maps (450, HDI_95%_ = [17, 6869]). After applying empirical criteria (see *Methods: HRF library generation*), the number of significant voxels reduced to 38,087 voxels from 110 maps of 76 scans (45, [5, 1662]). Applying temporal regularization, each estimated HRF time course showed smooth fluctuations, helping to improve the accuracy of HRF feature extraction (**Fig. 1a**).

**Fig. 1.**
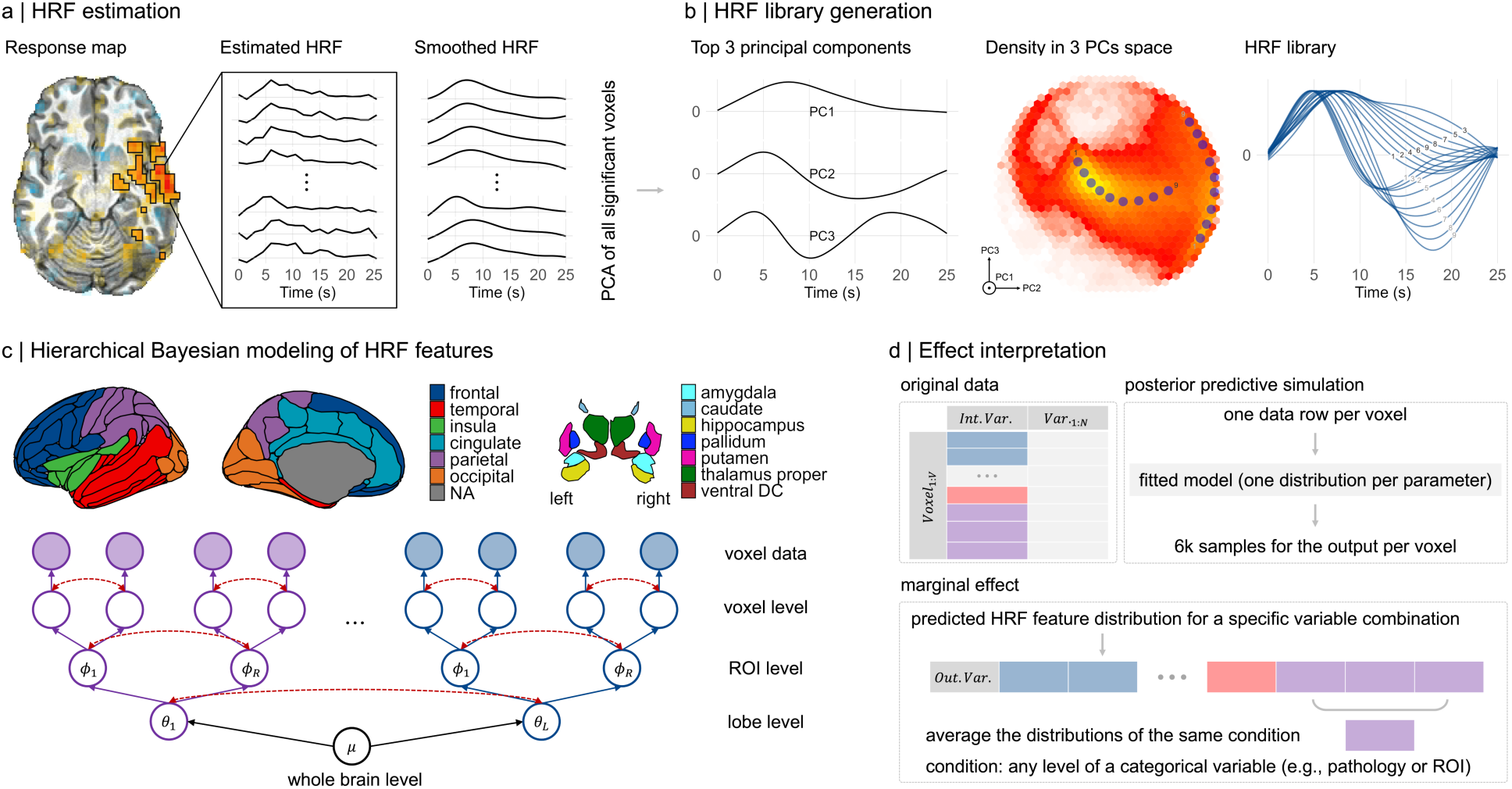
Overview of the methodological workflow. **a,** HRF estimation. *Left*: One fMRI scan from a patient typically results in a response map containing one or more significant BOLD clusters (black outlines) associated with epileptic discharges. *Middle*: HRFs (0–25 s; 17 time points) from significant clusters were estimated using AFNI’s deconvolution method using TNET basis functions. *Right*: Temporal regularization smoothed the HRFs to 4Hz to enable stable feature extraction and shape modeling. **b,** HRF library generation. *Left*: Principal component analysis (PCA) was applied to all significant HRFs across all response maps of all patients, resulting in the top three principal components (PCs) explaining 94% of the variance. *Middle*: For each HRF, the vector of three PC coefficients was normalized to unit length and projected into a 2D density map defined by PC2 (x-axis) and PC3 (y-axis), with PC1 pointing out of the plane (origin at the center). Each point represents a combination of PCs that reconstructs an HRF, with warmer colors indicating more frequent HRF shapes. There are two arcs tracing regions of high density. *Right*: Manually identified nine representative HRFs (blue dots) along each arc. **c,** A spatial hierarchical model (*i.e.,* parcel-lobe-brain) was constructed to partially pool HRF features within local parcellations, across lobes, and across pathologies. Models were estimated using Bayesian inference. **d,** Effects of interest were interpreted using posterior predictive simulations and marginal effects, defined as the influence of one variable (*e.g.,* pathology) on an HRF feature while holding others constant and averaging across all possible data values. Abbreviations: Int.Var., variable of interest; Out.Var., output variable. Var., other variables.

### HRF library featuring four shape groups in the epileptic brain (Study 1)

The variability in HRF shape can be sufficiently represented by three basis functions, derived from principal component analysis. The temporal decomposition method on all selected voxels resulted in the top three principal components (PCs) shown in **Fig. 1b**. In total, they explained 94.0% of the variance of all HRF time courses. PC1 captured a general trend of a typical HRF without an undershoot. PC2 and PC3 fluctuated faster than PC1 (*e.g.,* each passing the 0 baseline twice). When combined with PC1, their peaks before 10s can help modulate the HRF peak latency variation, and peaks after 15s can help model the HRF undershoot variation. The middle panel of **Fig. 1b** illustrates the density distribution of the three PCs coefficient combinations across all selected voxels. PC2 and PC3 define the x-y plane, with PC1 pointing orthogonally out of the page (origin at the center). Each point in this 2D density distribution plot corresponds to a specific combination of the three PCs that can reconstruct an HRF shape. More frequent combinations, representing more common HRF shapes, exhibit higher density (warmer colors). We visually identified two distinct arcs passing through the high density regions (blue dots in **Fig. 1b**) on the 2D density plot, resulting in 18 representative HRFs (right panel of **Fig. 1b**). We further grouped these HRFs into four groups based on their shape: Group 1 (no undershoot); Group 2 (low undershoot); Group 3 (moderate and early undershoot); and Group 4 (large and late undershoot; **Fig. 3d**). To compare with the canonical HRF, the average root mean square error (RMSE) to the canonical HRF is 0.55, 0.37, 0.23, and 0.56, respectively. These four shape groups provided a foundation for examining regional and pathological variations in HRF characteristics.

The resulting three PCs and the determined HRF library were robust when we varied the input voxels, indicating good generalizability for modeling HRF shape variability. Specifically, we tested different voxel selection strategies by varying the empirical ceiling threshold for HRF amplitude which controls the influence of potential spurious high amplitude voxels (10%, 5%, or 3%), and the upper limit of random subsampled voxels (k = 50, 100, 300, or 500; min[k, number of significant voxels] per IED related fMRI map, across 500 iterations). **Fig. 2a** shows all 6,000 PC sets (3 ceiling threshold × 4 subsampling × 500 random sampling), with darker colors representing higher ceiling thresholds. They exhibited stable overall fluctuation patterns with narrow variations. In all iterations, the resulting three PCs explained more than 93.0% of the total variance of selected voxels (**Fig. 2b**). There was only ∼1% increment in total variance explained when a less stringent ceiling threshold was used. The 2D density distribution plot of HRF shape (**Fig. 2c**) and the representative HRFs (blue dots) also remained stable compared to the main result shown **in Fig. 1c**, which used all significant voxels with a conservative 10% ceiling threshold.

**Fig. 2.**
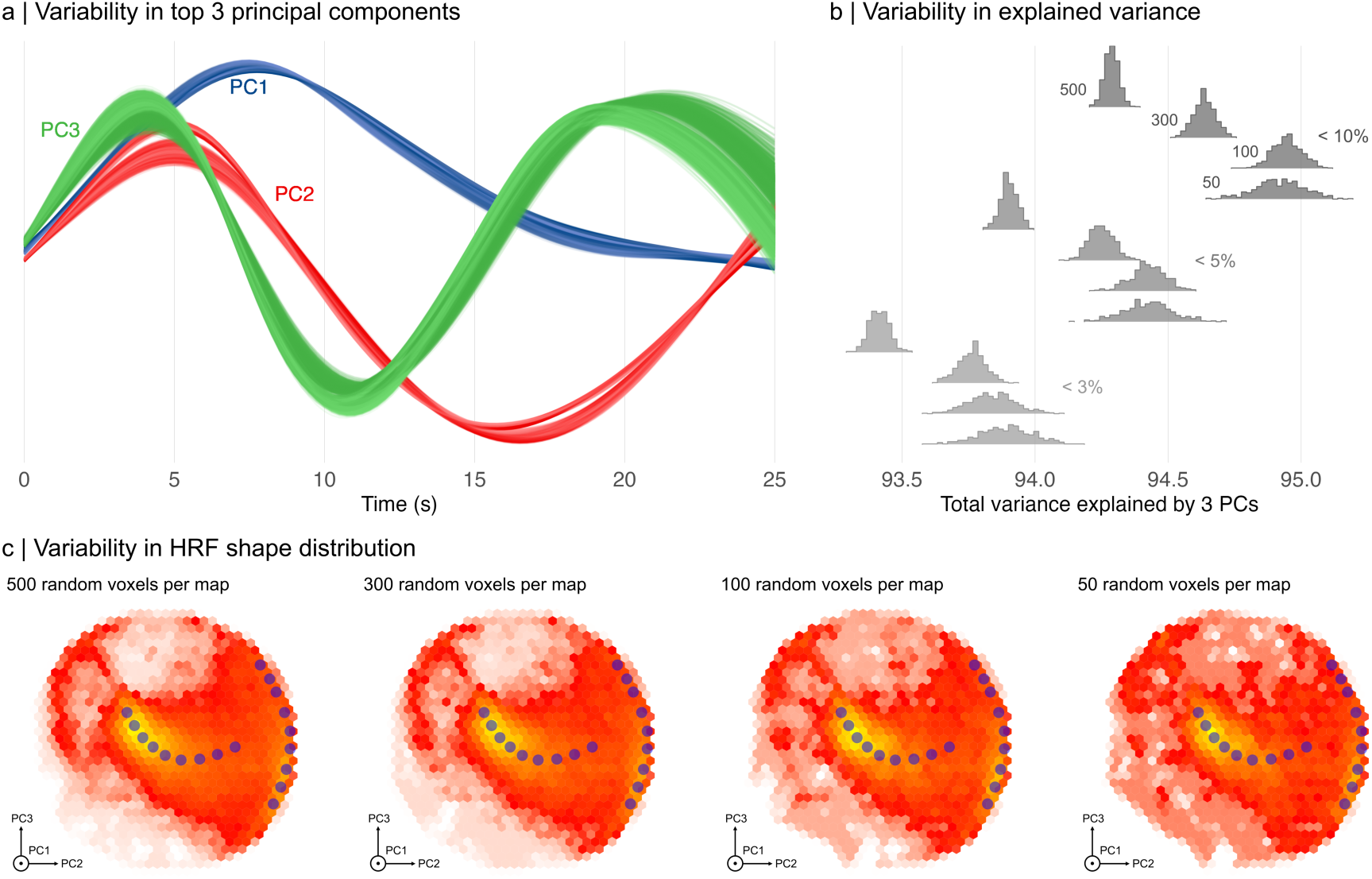
The top three PCs and HRF shape distributions are robust. **a,** Spaghetti plot of all 6,000 sets of the top three PCs obtained by varying the empirical ceiling threshold for HRF amplitude (10%, 5%, or 3%; darker colors indicate higher thresholds) and applying random voxel subsampling, *i.e.,* k = 50, 100, 300, or 500, min[k, number of significant voxels] per IED related fMRI map in each of 500 iterations. **b,** Histograms of total variance explained by the top three PCs across these 12 configurations (3 ceiling threshold × 4 subsampling). Bins are grey coded by ceiling threshold: 10%, 5%, and 3%, with subsample sizes increasing from bottom to top. All configurations explained at least 93.0% of the total HRF variance. **c,** 2D density distributions of HRF shapes using a 10% ceiling threshold and 50 to 500 random subsamples. PC2 and PC3 define the x-y plane, with PC1 pointing orthogonally out of the page (origin at the center). Density values are interpolated into hexagonal bins and normalized to the maximum for each subplot. Blue dots show the representative HRFs identified in the main results (*Fig. 1b*).

### HRF features show regional specificity in the epileptic brain (Study 2)

Having established the above HRF library, we next investigated whether HRF features vary systematically across brain regions. All selected voxels showed a strong match with the most correlated HRF from the library (maximum Pearson’s r = 0.76, [0.69, 0.99]). To avoid investigating the exact same voxels which also generated the library, a data-driven voxel selection was applied (see *Method: Modeling the associations*). This resulted in 18,881 voxels from 118 maps of 77 scans in the presumably non-pathological regions (*i.e.,* voxels outside the region subsequently resected). **Fig. S1a** shows the data distribution in this study. All parcels in the cortex and subcortical structures contained sufficient voxels (thresholded to have at least 5 voxels per parcel), with overall bilateral symmetry. The HRF features across these voxels were as follows: peak amplitude (0.2%, [0.0, 1.1]), time to peak (6.5s, [3.2, 10.2]), peak full-width at half-maximum (FWHM, 7.2s, [3.3, 12.5]), undershoot amplitude (-0.1%, [-0.7, -0.0]), time to undershoot (18.0s, [12.0, 24.2]), and undershoot FWHM (6.0s, [1.1, 11.8]).

HRF features exhibited a clear spatial organization following the brain’s anatomical hierarchy. We observed larger peak and undershoot amplitudes (**Fig. 3a-b**), along with earlier peak times (**Fig. S2a**), in primary cortices (*i.e.,* motor, somatosensory, auditory, and visual cortices) compared to other regions. Interestingly, this same pattern appeared in the spatial distribution of HRF shape group (**Fig. 3c**): Groups 3 and 4, characterized by moderate to large undershoot and early peak times (**Fig. 3d**), were mainly distributed in primary cortices. Additionally, Group 1, featured by no undershoot, was predominantly located in bilateral anterior cingulate regions. The remaining cortical areas mostly were in Group 2, which showed low undershoot amplitude. Among all features, both amplitudes and HRF shape group exhibited clear left-right symmetry in cortical and subcortical regions (**Fig. 3**). The symmetry was more consistent subcortically than cortically. For instance, time to undershoot and FWHMs were less symmetric in the temporal lobe (**Fig. S2**).

**Fig. 3.**
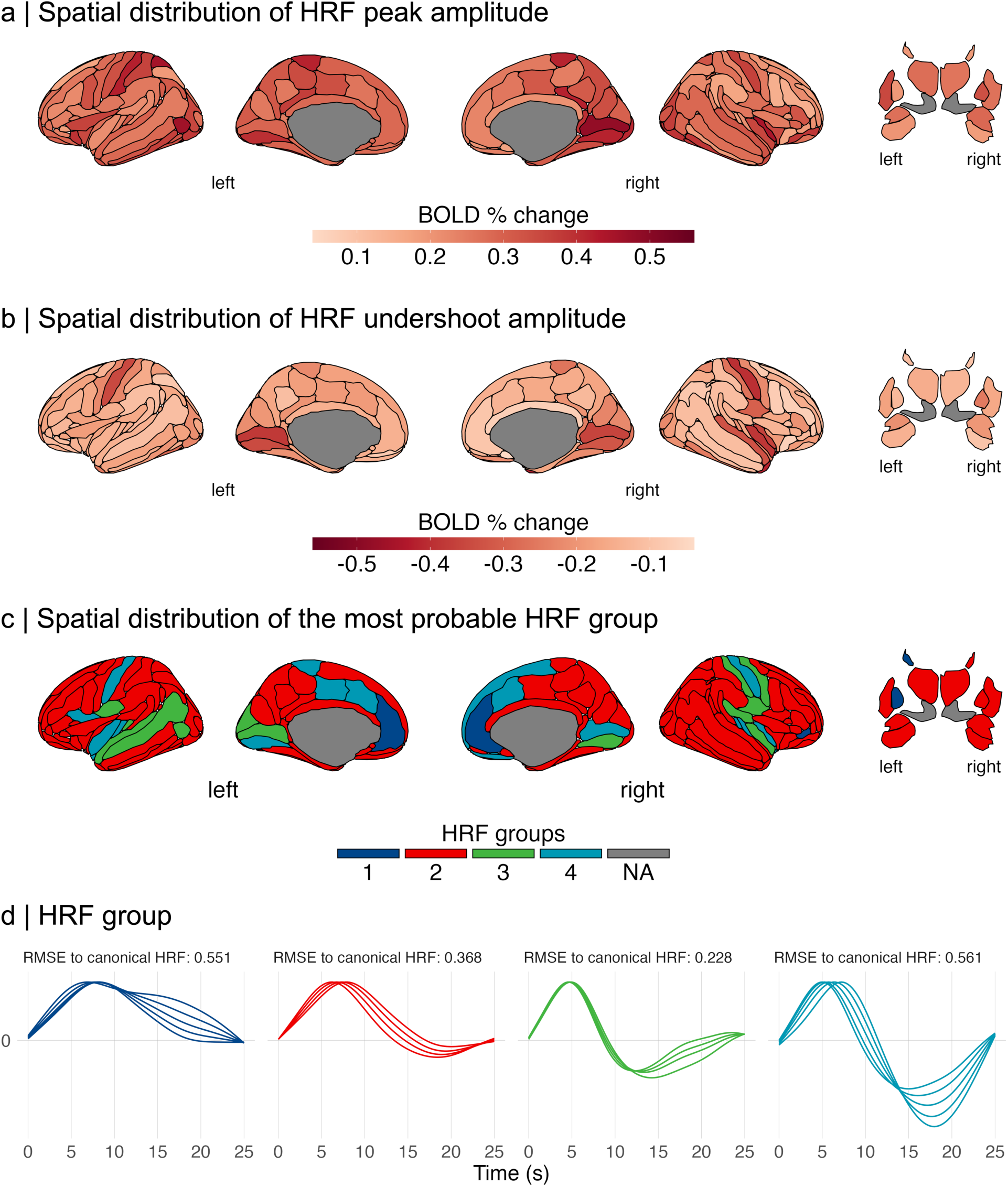
HRF features show anatomical location association in the epileptic brain. **a-b,** Parcel level spatial distributions of HRF peak and undershoot amplitudes. Analysis included 148 cortical parcels from the Destrieux atlas and 12 subcortical ROIs (derived from FreeSurfer). Parcels within primary cortices (*i.e.,* motor, somatosensory, auditory, and visual) exhibited larger magnitudes compared to other regions. **c,** Parcel level spatial distribution of the most probable HRF shape group (*i.e.,* the group with the highest marginal effect for each parcel). The pattern differentiating primary from other cortical areas observed in (a) is also shown here. **d,** HRFs in the library (*Fig. 1c*) were classified into four classes: Group 1 (no undershoot, blue), Group 2 (low undershoot, red), Group 3 (moderate and early undershoot, green), and Group 4 (large and late undershoot, cyan). To compare with the canonical HRF, the average root mean square error (RMSE) relative to the canonical HRF was 0.551, 0.368, 0.228, and 0.561, respectively. The canonical HRF parameters followed the *neuRosim* package defaults: peak at 6s, undershoot at 12s, dispersion of 0.9 for peak and undershoot, and an undershoot scale of 0.35.

### HRF features distinguish pathology types (Study 3)

We next examined whether HRF features also distinguish different pathologies (see *Methods: Dataset* for detailed pathology class definition). After significant voxel selection (see *Methods: Modeling the associations*), we identified 1,554 voxels from 48 maps across 38 scans within the surgical cavity. **Fig. S1b** shows the data distribution in this study. Parcels containing more than five voxels were primarily located outside eloquent cortices, with more voxels in the right hemisphere and within the bilaterally symmetric hippocampi.

All HRF feature distributions varied systematically across pathologies, indicating their clear associations. For instance, peak amplitude showed a three-level spectrum, increasing from HS/dual (*i.e.,* hippocampal sclerosis plus another pathology) and cortical malformation (*i.e.,* all types of focal cortical dysplasia) to normal (*i.e.,* no histopathological abnormalities were identified), and further to gliosis and encephalitis (**Fig. 4a**). The most probable HRF shape group followed the same trend (**Fig. 4a,c**) across pathologies, increasing from Group 1 in HS/dual, to Group 2 in cortical malformation and complex pathology (*i.e.,* mix of several pathologies), to Group 3 in normal, and finally to Groups 3-4 in gliosis and encephalitis. A similar pattern appeared in peak time (**Fig. S3c**) and FWHM (**Fig. S3e**), where normal showed the midrange values, separating lower values in gliosis and encephalitis from higher values in HS/dual pathology and cortical malformation. Undershoot features also reflected this pattern, particularly in amplitude (**Fig. S3b**). Undershoot time was lowest in normal appearing tissue, followed by gliosis and encephalitis, and highest in HS/dual pathology and cortical malformation (**Fig. S3d**). Undershoot FWHM increased from normal to gliosis and cortical malformation, then to HS/dual, and finally to encephalitis (**Fig. S3f**). Notably, HRFs in normal voxels showed the smallest difference to the canonical HRF typically used for healthy brains (RMSE = 0.23; **Fig. 3d**).

**Fig. 4.**
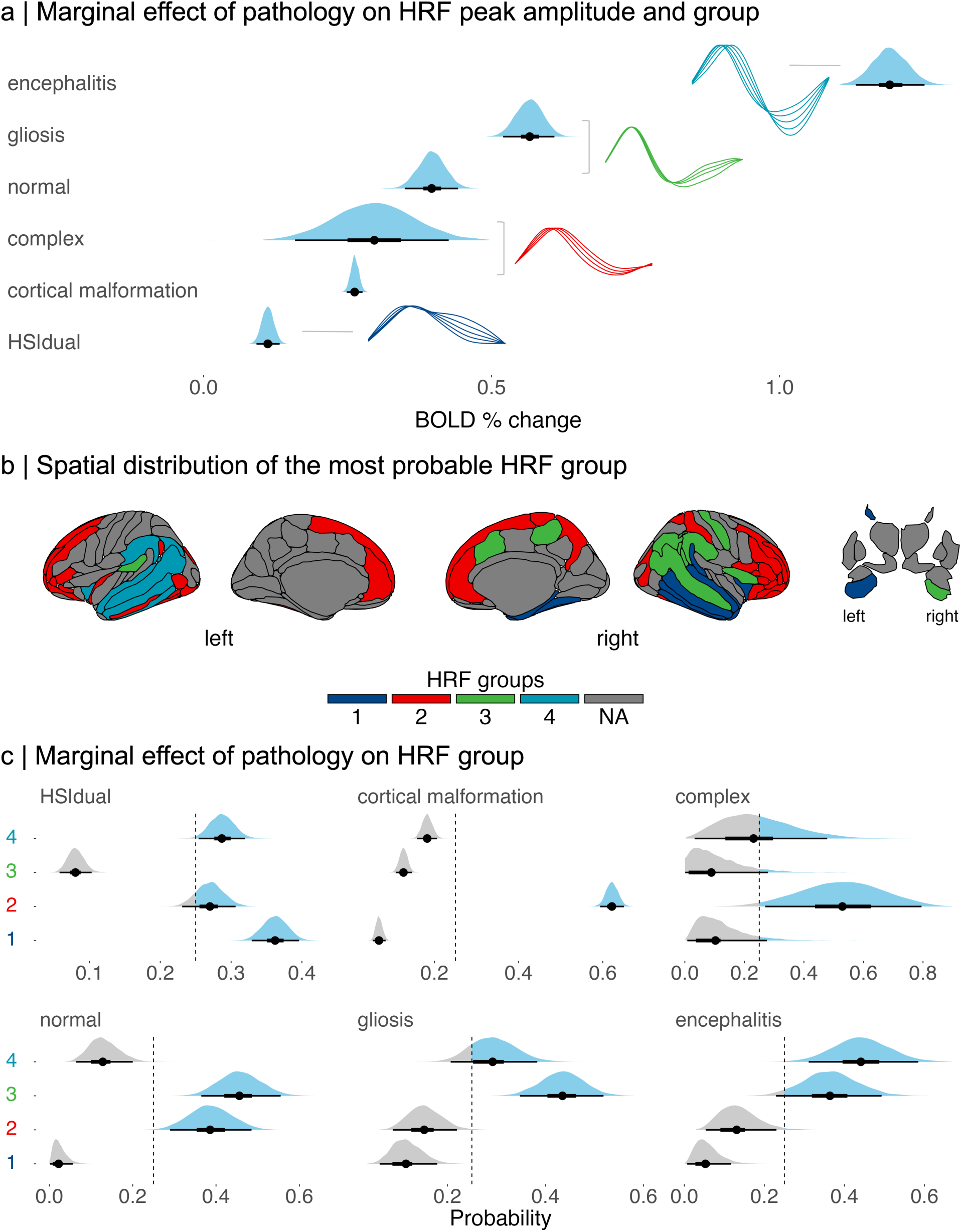
HRF features show association with pathology in the epileptic brain. **a,** Marginal effect of pathology on HRF peak amplitude and shape. The normal pathology type (*i.e.,* no histopathological abnormalities were identified) exhibited mid-range values, separating values observed in gliosis and encephalitis from those in HS/dual (*i.e.,* hippocampal sclerosis plus another pathology) and cortical malformation. For each pathology group, the HRF library and corresponding most probable HRF shape group (as determined in panel c) are shown, revealing a clear three level spectrum (HS/dual and cortical malformation ↔ normal ↔ gliosis and encephalitis). **b,** Parcel level spatial distribution of the most probable HRF shape group (*i.e.,* the group with the highest marginal effect per parcel). HRF shape groups showed pathology specific spatial distributions. **c,** Marginal effects of pathology on HRF shape group. Vertical dashed lines indicate the 25% chance probability across the four groups. Areas below this reference are shaded in gray. Each marginal effect distribution is shown as a blue density plot, with the black dot indicating the median, the thick black bar indicating the 50% highest density interval (HDI), and the thin black bar indicating the 95% HDI. Distribution widths illustrate the estimation uncertainty, *e.g.,* complex pathology (*i.e.,* mix of several pathologies) showed the widest distribution due to its smallest sample size.

Pathology accounts for additional HRF variability beyond that explained by anatomical location. Models that included both pathology and anatomical location (ranked first and second in **Fig. 5a**) achieved superior fit accuracy, generalizability, and parsimony compared to models excluding pathology (ranked third and fifth). The model including anatomical location and other factors but not pathology ranked third, whereas the model including pathology but excluding anatomical location ranked fourth. Clear contrasts in both HRF peak amplitude and HRF shape group were demonstrated between conditions with and without pathology in the same brain parcel (**Fig. S4**). Using HRF feature distributions estimated from data outside the surgical cavity (*Study 2*) as parcel-wise normative references, we found that the HRF peak amplitude distributions derived from data inside the cavity (with pathology; *Study 3*) were shifted toward the tails of the reference distributions for most parcels. The majority of parcels either changed their most probable HRF shape group or remained in the same group but with altered probabilities relative to the corresponding normative reference.

**Fig. 5.**
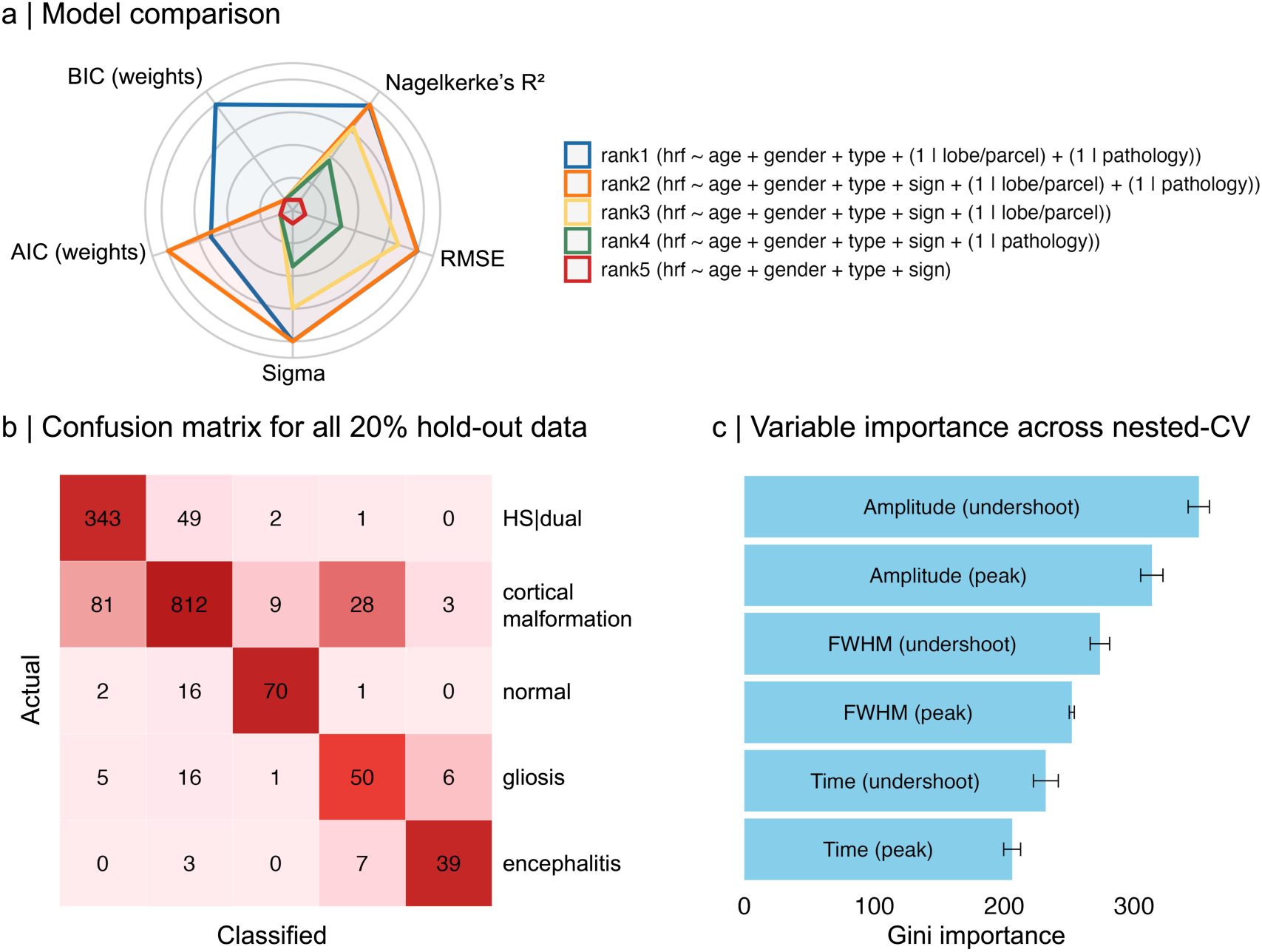
Associations between HRF features and pathology revealed by model comparison and classification. **a,** A radar plot shows model comparison results across five candidate models, ranging from a baseline fixed effects only model to a full model including all fixed and random effects. Each axis represents one model performance metric: Nagelkerke’s pseudo-R², root mean squared error (RMSE), Akaike information criterion (AIC), and Bayesian information criterion (BIC). Each metric was normalized and rescaled from 0% to 100% relative to the best performance across models. Models were ranked by overall performance across all metrics. **b,** Confusion matrix from pathology classification using all 20% hold-out data of the 5×5 nested cross validation. The highest values appear along the diagonal, indicating accurate classification. **c,** Gini importance of HRF features from the classification model. Feature importance decreased from amplitude to FWHM and then to time, with undershoot related measures showing slightly higher values within each category. Black error bars indicate the standard deviation across cross validation folds. *hrf:* HRF shape group; *type*: IED type; *sign*: HRF sign.

Given the above estimated effects of pathology on HRF features, we further demonstrated these associations by classifying pathology using only six HRF features. Testing on hold-out data within a 5×5 nested cross-validation framework showed good performance (precision = 0.79 ± 0.04, recall = 0.79 ± 0.05, F1 = 0.79 ± 0.04, MCC = 0.74 ± 0.03, and Cohen’s κ = 0.74 ± 0.03). The confusion matrix on the hold-out data showed the highest values along the diagonal, confirming accurate classification (**Fig. 5b**). Feature importance, measured by Gini importance, decreased from amplitude to FWHM and then to time, with undershoot related measures showing slightly higher values within each category. All performance metrics were significantly higher than their corresponding null distributions derived from 1,000 random permutations of the pathology labels (**Fig. S5**). Finally, when HRF features were used to classify cortical lobes instead of pathology, the overall performance metrics dropped by around 0.2 (precision = 0.62 ± 0.06, recall = 0.55 ± 0.06, F1 = 0.57 ± 0.05, MCC = 0.56 ± 0.05, and Cohen’s κ = 0.56 ± 0.05).

### Associations remain robust to a different parcellation atlas

We also replicated Studies 2 and 3 using the Schaefer2018 atlas (200 parcels), which integrates local and global intrinsic functional information^37^, combined with the subcortical parcellation from FreeSurfer. The results were consistent with those shown above. We observed a small increase in spatial specificity due to the smaller parcel size (200 vs. 148), while the overall spatial distribution of HRF features remained largely unchanged. **Fig. S6** reproduces the associations between HRF features and anatomical location from *Study 2* (**Fig. 3**), and **Fig. S7** reproduces the associations between HRF features and pathology from *Study 3* (**Fig. 4**). **Fig. S8** presents the corresponding model comparison and pathology classification results from **Fig. 5**. Moreover, pathology classification performance is similar to the main analysis (precision = 0.82 ± 0.02, recall = 0.79 ± 0.02, F1 = 0.80 ± 0.02, MCC = 0.75 ± 0.02, and Cohen’s κ = 0.75 ± 0.02).

### Effects of other factors on HRF are subtle

The spatial distributions of HRF shape groups resembled the organization of primary vs. other cortices previously shown in **Fig. 3c** when stratified by positive (activation) and negative (deactivation) HRFs in **Fig. S9a-b**. Positive HRFs mainly fell into Group 2 (41.7% [40.9-42.6]) and Group 4 (29.8% [29.1-30.6]; **Fig. S9c**), with the remaining voxels evenly divided between Groups 1 and 3. Negative HRFs were similarly distributed across Groups 1-3, with a low probability of being Group 4 (9.6% [8.9-10.4]; **Fig. S9c**).

Overall, other factors had only minor effects on HRF group. The absolute probability changes of belonging to a given HRF group with a 5-year increase in age was less than 4%, with the largest effect observed in Group 1 (-3.2%, [-3.4, 2.9]; **Fig. S10a**). Both females and males showed the highest probability for Group 2 (F: 34.8%, [33.9, 35.7]; M: 41.9%, [40.8, 42.8]; **Fig. S10b**). The difference was mainly in the secondary groups where males showed a higher probability for Group 4 (28.9%, [28.0, 29.7]), and females showed a higher probability for Group 3 (26.6%, [25.6, 27.4]). Similarly, both long-lasting IEDs and spikes were most frequently associated with Group 2 (37.8%, [37.1, 38.5] and 40.1%, [38.1, 41.9], respectively), with the other three groups each accounting for approximately 20% (**Fig. S10c**).

## Discussion

This study provides the first systematic investigation of HRF variability across the entire epileptic brain, and its modulation by anatomical location as well as pathology, using a large and well-characterized EEG-fMRI dataset. We identified 18 representative HRF shapes that grouped into four distinct shape groups and further mapped their spatial distribution throughout cortical and subcortical regions, revealing structured rather than random variability. Furthermore, we estimated the distribution of HRF features across different pathologies and demonstrated their association using effect estimation, model comparison and classification. These findings suggest that HRF shape is modulated by anatomical location and pathology, and emphasize the limitations of traditional approaches based on canonical or multiple predefined HRFs. This study establishes a foundation for HRF modeling of EEG-fMRI analysis in epilepsy. These insights may refine presurgical evaluation using EEG-fMRI, improve localization of epileptogenic zones, and inform broader understanding of neurovascular dynamics in neurological disease.

### HRF temporal variability in the epileptic brain

By originally combining two advanced methods, regularized HRF deconvolution^18^ and temporal decomposition^34^, we addressed the question raised in the introduction: how temporally variable is the HRF in the epileptic brain. Across both voxel level summaries (*Study 1*) and parcel level inferences (**Fig. 3ab** and **S2**), we observed substantial deviations from patterns reported in healthy brains. Compared to healthy subjects^16,17,21^, HRFs in epileptic patients showed a small delay in peak timing (mean 6.6s vs. 4.6-6.2s), with a notably larger standard deviation (SD 1.9s vs. 0.4-0.8s), wider peaks (mean±SD 7.5±2.4s vs. 4.1-4.5±0.5-0.7s), and considerably later undershoots (mean 17.0s vs. 10.8-13.6s). These findings align with our previous work showing wider, delayed HRFs during auditory motor tasks in epilepsy^32^ and earlier observations of wide peak latency ranges (2.5-9.5s), and delayed undershoots (∼21.2s) in smaller (N=15) epilepsy cohorts^38^.

While summarizing individual HRF features can be informative, these values are inherently correlated^16,17,21^ by the shape *per se*, making isolated interpretations confounded by their mutual dependencies. Our HRF library instead naturally integrates these correlated features into representative patterns, allowing a compact and intuitive description of hemodynamic variability and provides an estimate on how many representative HRFs can be identified in the epileptic brain. Remarkably, among all possible HRF variations, our data-driven HRF library uncovered a rich yet structured spectrum of hemodynamic profiles rather than random fluctuations. We identified 18 representative HRF shapes that categorized into four shape groups. The aforementioned wider peak (Groups 1 and 2) and delayed undershoot (Groups 2 and 4) are also reflected in our resulting HRF library. The library naturally captures feature correlations reported in healthy cohorts^16^: the positive relationship between peak and undershoot timing (they shift together following the order of Groups 3, 4 and 2), and earlier peaks are associated to narrower widths (Groups 3-4 vs. 1-2).

These epilepsy specific HRF shapes may serve as data informed alternatives to the multiple shifted gamma functions currently used in many epilepsy studies^4,27,28^, which typically vary only by peak timing (3-9s) and which lack realistic undershoots. In contrast, Groups 3-4 exhibit moderate to large undershoots, consistent with previous reports showing undershoot over peak ratios of 0.54 (0.02-1.21)^38^. The HRFs without undershoot in Group 1 may also help better capture the negative BOLD response, as this shape appeared more frequently among negative HRFs localized within the default mode network (**Fig. S9b**), regions typically associated with IED related deactivations^39^. Similar HRF shapes for negative responses have been reported in task-based studies of healthy participants^16,17^ and in epilepsy^38^, where negative responses were found to differ from the simple mirror of the canonical HRF.

Our approach leverages PCA to summarize HRF variability efficiently. PCA has long been used to characterize hemodynamic response^40^, hypothetical^41,42^ or empirical^34,43^ HRFs derived from healthy participants. Consistent with prior work in healthy individuals^43^, the top three components in our data explained sufficient total variance (94%) across all HRFs, with results stable under different voxel selection thresholds and subsampling (**Fig. 2**). This robustness validates PCA as a reliable framework for quantifying HRF variability in epilepsy. Our PCA basis and HRF library may provide a more physiologically grounded and reliable framework for mapping IED related responses: researchers can incorporate the top three PCs as basis functions to capture epilepsy specific hemodynamics, constrain their combinations^42^ if necessary, or directly replace traditional HRF sets^27,28^. Similar empirically derived HRF sets have improved single trial effect estimation and image decoding in healthy participants^29^, opening new avenues to explore IED level dynamics and associated decoding in the epileptic brain.

### HRF features vary across space

Beyond temporal diversity, HRF features exhibited a clear spatial organization following the brain’s anatomical hierarchy. HRF amplitudes and shape groups showed a systematic and largely bilateral pattern. Primary cortices including motor, somatosensory, auditory, and visual regions exhibited larger peak and undershoot amplitudes (**Fig. 3a,b**), earlier peak latencies (**Fig. S2a**), and a preference of Groups 3 and 4 (**Fig. 3c**), characterized by moderate to large undershoots and slightly faster dynamics. This organization is consistent with observations in recent studies of healthy subjects^16,18,21^, suggesting that HRF spatial distribution are preserved in presumably non-epileptogenic regions of the epileptic brain.

Although most patients in our cohort had unilateral IEDs, where lateralized and regional IED related responses are expected, we still observed a systematic, bilateral spatial organization of HRF features. Similar topographies have been reported in task based studies of healthy individuals^16–18,21^. However, these findings cannot fully rule out potential bias from the inherent bilateral nature of task evoked responses. The convergence between our results and those from healthy cohorts, despite fundamentally different sources of neural activation, suggests that HRF spatial organization may be intrinsically associated with anatomical location and functional properties rather than being context dependent. Furthermore, at the group level by aggregating activated parcels across both cortical and subcortical regions, our analysis achieved whole brain coverage exceeding the 68-90% coverage engaged by multiple tasks in healthy fMRI studies^16–18,21^. We acknowledge, however, that sample sizes per voxel remain limited due to the heterogeneous nature of IED responses across patients. To address this, we developed a spatial hierarchical model (**Fig. 1c**) that performs partial pooling of HRF features across parcels, lobes and brain. This approach allows local estimates to be informed by both regional and global homogeneity, providing statistically demonstrated improvements in accuracy, reliability, and generalizability over conventional voxel wise inference^44,45^.

The spatial structure of HRF features aligns with the principal functional gradient of the brain, which extends from unimodal primary sensory and motor areas to transmodal association cortices^46,47^. In numerous previous studies^48–51^, this gradient has been shown to recapitulate the primate cortical hierarchy, and also serves as a potential axis of intrinsic functional dynamics, with longer neuronal time scales being found towards its transmodal apex. Indeed, sensory and unimodal regions exhibited earlier and larger peak (Groups 3 and 4), whereas transmodal cortices showed relatively weaker, delayed responses (Groups 1 and 2). Unimodal regions are known for their dense intracortical myelination, which declines along the sensory to association axis^52,53^, and for systematic variation in excitation-inhibition balance that modulates how strongly structural connectivity constrains functional dynamics^54^. These properties may support the similar spatial distribution of HRF observed here. Because the HRF mediates the transition from neural activity to BOLD signal, its spatial organization likely reflects underlying neurovascular mechanisms that contribute to the macroscale gradients observed in resting state connectivity. A promising next step would be to integrate the HRF spatial map (*e.g.,* amplitude and shape) from this study with intracranial stereoelectroencephalography (SEEG)-based cortical atlas that capture normative electrophysiological activity across regions^55^, or to directly leveraging simultaneous SEEG-fMRI data^56,57^. For instance, our previous work using such a normative SEEG atlas showed that gradients derived from electrophysiological activity reflect the similar unimodal-transmodal cortical organization observed in fMRI^58^. By convolving region-specific electrophysiological signals with their corresponding HRFs estimated in the current study, future work could test whether neurovascular coupling contribute to the macroscale functional gradients seen in fMRI.

### HRF features are modulated by epileptogenic pathology

One advance of this work is the discovery that HRF features systematically differ across pathology types, revealing a potential hemodynamic signature of underlying tissue pathology. An intuitive way to interpret these results is to jointly consider HRF peak amplitude and shape group. Since HRF shapes are normalized by amplitude, this combination effectively reconstructs the HRF profile. They all followed a three level pathology spectrum (HS/dual and cortical malformation ↔ normal ↔ gliosis and encephalitis; **Fig. 4a**): with peak amplitude and group index increasing from HS/dual and cortical malformation (Group 1-2), through normal tissue (Group 3), to gliosis and encephalitis (Group 3-4). The spatial distribution of the most probable HRF shape groups closely matched the known spatial distribution of the pathologies^59–61^ that preferred those same HRF shape groups. In contrast, when HRFs were estimated from non-pathological data (outside the cavity) for the same parcels, parcels either shifted to a different preferred HRF shape group or showed altered probabilities (**Fig. S4**).

Although the physiological explanation remains challenging, we demonstrated the association between HRF features and pathology also using model comparison and machine learning classification. Specifically, our results showed that pathology modulates the HRF in addition to the underlying effects of anatomical location. Models incorporating pathology and anatomical location achieved the best overall performance (**Fig. 5a**). The model including anatomical location but not pathology ranked above the model including pathology but excluding location. Together, these findings indicate that pathology provides additional explanatory power to location. We also observed clear contrasts in both HRF peak amplitude and HRF shape group between conditions with and without pathology in the same brain parcel (**Fig. S4**). Moreover, HRF features alone classified pathology with good accuracy. Although this performance may be partially inflated by correlations between pathology and location, it still outperformed the classifications on lobes and networks, with a nontrivial overall performance difference of ∼0.2. One limitation is the imbalance in sample size across pathology groups (**Fig. S1b**), which we addressed through weighting and resampling in the classification. We note that the pathology classification was designed to demonstrate associations between HRF features and pathology. Larger datasets will be required to assess the extent to which HRF features alone can support pathology classification. In practice, combining HRF features with other anatomical microstructure features may provide a more robust approach to non-invasive pathology estimation.

Our results provide the first evidence linking HRF features to pathology in epilepsy. In our previous study, no significant differences in HRF peak amplitude or latency were found at the maximum *t*-value voxel between patients with cortical malformation and those with HS^33^. However, the maximum *t*-value voxel does not necessarily locate in the pathological region. In this work, we constrained significant voxels to the surgical cavity, ensuring closer correspondence to pathology. Another epilepsy study suggested that noncanonical HRFs were likely artifacts rather than true pathological responses^62^. Our stringent statistical and empirical filtering (see *Methods: HRF library generation*) minimizes such artifacts, and the fact that all four HRF shape groups observed here have been reported in healthy brains further supports their validity. To our knowledge, no prior studies have classified epilepsy pathology using HRF features. One study distinguished subjective cognitive decline from healthy controls with a 94.1% accuracy^30^. Although our accuracy is lower, classifying multiple pathology types is a more challenging task.

### Effects of other factors on HRF and limitations

One large healthy cohort study did find significant HRF feature differences between young and old groups (mean age 25 vs. 64), but the effect size is subtle^19^. We did not observe considerable age effects, likely because the oldest participant was younger than 60. Moreover, even if our work was based on a relatively large cohort of well-characterized surgical candidates, addressing developmental and lifespan changes likely requires inclusion of additional datasets, which may motivate follow-up studies in multi-center cohorts. One limitation of this study is that pathology outside the surgical cavity remains uncertain. Although our assumption that tissue outside the cavity is relatively less pathological compared to the cavity itself is overall reasonable, IEDs related fMRI primary clusters do not always predict surgical success (individual level accuracy 0.60-0.66)^6^, and overall seizure freedom rates after surgery remain around 64%^2^. These findings, together with post-mortem findings^63,64^ and a rich MRI-based literature suggesting large-scale compromise in traditional focal epilepsy syndromes^65–69^, indicates that residual pathology may exist beyond resected regions. This uncertainty also reduced the sample size for *Study 3*. One possible way to address this limitation is to increase the number of pathological samples using clear anatomical imaging informed pathological regions, though such clear cases are often not considered for presurgical evaluation with other modalities.

The physiological basis of the BOLD signal remains incompletely understood, making the explanation of our results challenging. Here, we propose a hypothesis to account for the pathology specific HRF differences we observed. Tissue in the case of HS/dual and cortical malformation is typically firm, whereas gliosis and encephalitis are softer^70–73^. Following the BOLD balloon model^13,74–76^, we use tissue stiffness as a proxy for venous compliance. Softer, more compliant tissue would delay cerebral blood volume recovery and thereby increase the BOLD undershoot, whereas stiffer tissue would allow cerebral blood volume to recover rapidly and reduce undershoot. This relationship between venous compliance and undershoot amplitude has been demonstrated in a simulation study^77^. In addition, gliosis is associated with astrocytic hypertrophy and proliferation, and astrocytes play a key role in neurovascular coupling^78,79^. It is also a reactive response to injury that likely elevates metabolic demand, contributing to the increased HRF amplitudes. A 7T study of mesial temporal lobe epilepsy reported greater asymmetry in hippocampal vessel density compared with controls, and reduced ipsilateral vascular density to the presumed seizure onset zone^80^. This reduction may explain the lower HRF amplitudes observed in HS/dual pathology. However, as counterexamples, increased vascular density was reported within the focal cortical dysplasia^81,82^. Together, our findings suggest that both location and possible pathology induced microstructural changes jointly shape HRF variability in the epileptic brain. Further integration of electrophysiological, vascular and hemodynamic measurements may provide deeper insight into how anatomical and pathological factors interact to modulate neurovascular coupling in epilepsy.

## Methods

### Dataset

We analyzed an EEG-fMRI dataset collected at the Montreal Neurological Institute and Hospital (MNI). Prior to participation, every patient signed a written consent form (approved by the Research Ethics Committee of the MNI). The dataset contains 84 scans from 79 patients (49% female; median age: 26 years, range: 11-53, 5 patients had 2 scans). After the EEG-fMRI scan, each patient underwent one or more surgical resections (101 in total, range: 1-3), all of which included a minimum one year follow-up with postoperative brain imaging. Surgeries targeted the temporal lobe (57%), frontal lobe (28%), parietal lobe (6%), occipital lobe (3%), and multiple lobes (6%). Experienced epileptologists (A.K. and T.A.) classified the pathology for each surgical resection based on the histopathology report, following established criteria^83–86^. Pathologies were categorized as: cortical malformation (40%, including all types of focal cortical dysplasia); hippocampal sclerosis and dual pathology (21%, HS/dual, dual pathology is defined as HS plus another pathology, e.g., HS with focal cortical dysplasia); gliosis (21%); normal (6%, no histopathological abnormalities were identified in the specimen); encephalitis, tumors, or vascular malformations (4%); complex pathologies (2%, e.g., hemimegalencephaly and focal cortical dysplasia with balloon cells; hamartomas, dermoid cyst and gliosis); and cases with unavailable histopathology analysis (6%). Given that histopathology provides a region level diagnosis, we assigned a uniform pathology label from the above groups to each surgical cavity for subsequent analyses.

### EEG-fMRI scan protocol

All neuroimaging data were acquired in a 3T Siemens MR scanner (Siemens, Germany). The protocol for each patient included a high-resolution T1-weighted anatomical scan (1.0 mm isotropic) followed by a series of T2*-weighted functional runs. Each fMRI run lasted 6 minutes, resulting in a total acquisition time of 60-90 minutes. We used two sets of fMRI parameters: 82% of scans were acquired with a repetition time (TR) of 1.90s and 3.7mm isotropic voxels, whereas the remaining 18% used a TR of 1.75s and 5.0mm isotropic voxels. We recorded concurrent scalp EEG inside the scanner and synchronized it with the fMRI data using an MR-compatible BrainAmp EEG system (Brain Products GmbH, Germany). The EEG montage followed the 10-20 system but incorporated six additional electrodes from the 10-10 system (F9/10, T9/10 and P9/10) for enhanced coverage of the temporal regions.

### EEG data processing

We processed the EEG data in BrainVision Analyzer (Brain Products GmbH, Germany) to correct for MR gradient and ballistocardiogram artifacts using the average artifact subtraction (Allen et al., 1998, 2000). Experienced epileptologists (A.K. and H.M.K.) manually annotated the onset and duration of each IED. Based on clinical information, IEDs that assumed to originate from a common brain region were grouped and referred to as a single IED type. Each IED type was modeled as a separate event in the fMRI analysis and is referred to as fMRI map hereafter. We simply classified IED types into two general categories based on their duration: brief (*e.g.,* spikes) and long-lasting (*e.g.,* bursts or polyspikes) discharges.

### Anatomical and functional image processing

Anatomical and functional images of EEG-fMRI acquisitions were arranged following the Brain Imaging Data Structure^87^. Individual preoperative T1-weighted MRI processing included: intensity non-uniformity correction and skull-stripping using *@SSwarper* of AFNI^88^; followed by cortical and subcortical segmentations using *recon-all* of FreeSurfer^89^. The postoperative images (T1-weighted MRI or CT images) were coregistered to patient specific preoperative images. The surgical resection cavity mask was initially defined using a semiautomatic tool^90^, and an experienced neurophysiologist (C.A) then visually validated and refined it if necessary.

fMRI data were preprocessed in individual space using *afni_proc.py* of AFNI^88^. Preprocessing steps included: removing the initial non-steady-state volumes; correcting voxel wise BOLD time course outliers (despike); slice timing correction; aligning BOLD volumes to the volume with minimum outliers; applying rigid body functional to anatomical registration; masking functional data within the brain mask; spatially smoothing the data within the brain mask to 6.5mm full-width at half-maximum (FWHM) in all three axes, which accounts for differences in voxel size across the dataset; and scaling voxel-wise data by the local average which allows the regression coefficients to be approximately interpreted as percent signal change^91^. We censored BOLD volumes when Euclidean norm of the motion parameters is larger 1.0mm, or more than 5% of the brain voxels were detected as outliers. We also resampled tissue segmentation and the surgical cavity mask to the BOLD spatial resolution for later feature extraction. Finally, we performed quality control using an automatically generated QC report^92^.

### HRF estimation

We estimated the voxel-level HRF in two steps: deconvolution and temporal regularization (**Fig. 1a**). We performed deconvolution using *3dREMLfit* of AFNI on concatenated BOLD runs using a GLM with the following components: run wise baseline, drifts, and 6 motion regressors; voxel-wise 17 AFNI’s TENT basis functions from 0.0 to 25.5s; and voxel-wise ARMA(1,1) modeled autocorrelation in the residuals. GLM was solved using restricted maximum likelihood estimation providing *F*-statistic of the model as well as per impulse response function *t-value*. We then applied temporal regularization using spline smoothing^18^ on the resulting HRF to reduce overfitting and increase the temporal resolution. This process resulted in an HRF at a resolution of 0.25s for later feature extraction and HRF shape modeling.

### HRF library generation

A library was generated to represent the typical HRF shapes estimated from the entire dataset following two steps: stringent voxel selection, keeping only voxels containing statistically and practically significant HRF estimations; and temporal decomposition, which summarizes typical HRF shapes from all selected voxels.

We first selected voxels based on statistical significance, using a two-sided *p*-value threshold of 0.001 for both the GLM model *F*-statistic and the maximum *t*-value across all 17 weights associated with the TENT basis functions (*i.e.,* every 1.5s from 0.0 to 25.5s). Then, to ensure the HRF was practically meaningful, we considered four empirical criteria: the HRF magnitude did not exceed 10% signal change considering the ceiling BOLD percentage change in 3T^93^; peaks occurred within 2-10s; the HRF started close to 0 (*i.e.,* amplitude at 0s stayed within ±0.25% signal change); and returned to baseline after 20 s (*i.e.,* at least one value after 20s stayed within ±0.25% signal change). Negative HRFs were flipped to only consider the shape. Finally, only HRF within the grey matter mask were considered.

We then applied the temporal decomposition method^34^ on the selected voxels (**Fig. 1b**) to extract typical HRFs. We first performed PCA on all HRF time courses and selected the top three principal components (PCs). Each HRF was then represented by its corresponding three PCs coefficients after normalizing the coefficients vector to unit length. The PC coefficients vectors, representing specific HRF shapes, were projected onto a 2D space (two PCs in the x-y plane) to create a visualization of HRF shape density distribution. Finally, we manually defined two arcs (manifolds) representing the high density trajectories using nine points each on the density plot.

We also assessed the robustness of the resulting three PCs and typical HRF shapes using a random subsampling procedure on the significant voxels from each IED-fMRI map. We first considered the other two values, 5% and 3%, for the first empirical criterion regarding the BOLD ceiling percentage change. Then, in each of 500 iterations per map, we sampled min[k, N] voxels, where N is the total number of significant voxels and k is a predefined threshold (50, 100, 300, or 500). Subsequently, for each iteration and threshold, we compared the variance explained by the top three PCs and their corresponding time courses, as well as the HRF shape distribution plot.

### Modeling the associations

We then studied the association between HRF characteristics and anatomical location and pathology following two steps: HRF feature extraction; and effect modeling, which models the effect of variables of interest on the HRF characteristics.

To avoid the mixture of positive and negative HRFs in the effect modeling, we reversed negative HRFs, defined as those with a negative peak within 2 to 10s and a negative area under the entire HRF curve. The original sign was saved as a binary HRF feature (positive or negative) to study the effect of HRF sign. Six HRF features were then extracted using the MATLAB *findpeaks* function, including amplitude, latency and width (full width at half maximum, FWHM) for both peak and undershoot. To study HRF shape, we assigned each HRF to one of the 18 typical HRF shapes in the library exhibiting the highest Pearson’s correlation. Considering the balance of model complexity and sample size, this label was defined after grouping the 18 typical HRF shapes in the library into four groups sharing similar shape (**Fig. 3d**). The voxels used here were designed not to be the same as the voxels used in the HRF library generating section to avoid overfitting, that is, to avoid investigating the exact same voxels which generated the library. Therefore, we used a different voxel selection criterion: statistical significance, HRF amplitude did not exceed 10% signal change, correlation values to the HRF library larger than 0.7 (*i.e.,* the lower bound of the HDI_95%_ of all correlation values), and the amplitudes of the peak and undershoot were smaller than 2.5%, with a peak latency within 3 to 13s (thresholds defined based on the feature value distribution). To avoid including less representative parcels and ensuring a minimum sample size for later parcel level pooling, we further filtered out parcels that had less than 5 selected voxels.

The resulting HRFs were then used to study two main questions: the effect of anatomical location on HRF features (*Study 2*) and the effect of pathology on HRF features in addition to the location effect (*Study 3*). In *Study 2*, we considered voxels outside the region subsequently removed (referred to here as the “surgical cavity”) with abnormal histopathology (*i.e.,* pathological cavity), referred to as the presumably non-pathological regions. In *Study 3*, we only included voxels inside the surgical cavity.

To study the effects of anatomical location and pathology on the HRF features, we proposed a novel spatial hierarchical model (**Fig. 1c**). This model, extends from our previous work^94^, uses partial pooling of HRF features within local parcellations, across lobes, and across pathologies. It allows the effects to be estimated not only by voxel data but also adjusted according to the local and global homogeneity of the effect, resulting in more generalizable estimation. The anatomical hierarchical structure considered the levels of parcel, lobe, and whole brain, whereas the pathology hierarchical structure only considered one level. We also involved other factors, including age, gender, IED type (spike or long-lasting) and HRF sign (positive or negative). In general, the simplified model notation is as follows,

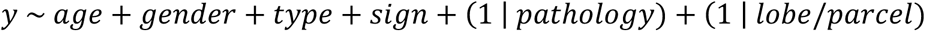

where *y* is the HRF feature of one voxel following a likelihood function. When the HRF feature is shape group, the likelihood function is a categorical distribution (multinomial distribution); whereas it is a normal distribution when the HRF feature is continuous (*e.g.,* peak amplitude). The notion of | represents the hierarchical structure, and / represents the nested multi-level hierarchical structure. We assigned weakly informative priors for all parameters (see code for details).

We considered 148 parcels from the Destrieux atlas^95^ and 12 subcortical ROIs from Freesurfer segmentations (**Fig. 1c**). These were grouped into 12 cortical lobes (left and right temporal, frontal, insula, parietal, cingulate, and occipital) and left and right subcortical regions. Given that effects on HRF in the epileptic brain may or may not be symmetric, we separated the left and right hemispheres.

### Statistical inference and effect estimation

We used Bayesian inference to estimate probability distributions for all model parameters. Unlike traditional methods that produce a point estimation with uncertainty range, Bayesian inference describes the estimated parameter and uncertainty with a probability distribution, leading to more accurate, reliable, and generalizable estimation^96,97^. In simple terms, Bayesian inference repeatedly samples different combinations of parameter values and checks how well each combination fits the data using Bayes’ theorem. After many iterations, these samples form a distribution that shows how likely each parameter value is, given the data. We estimated all parameters together using the Hamiltonian Monte Carlo algorithm^98,99^ implemented in the R package *brms*^100^. For the parameter of each model, the posterior distribution was sampled by 6,000 samples. Diagnostic checks were performed to ensure the model fit was reliable^101^. Models were run on an M4 Pro CPU, sampling 6000 samples per parameter in under 30 minutes.

After the model fitting, we estimated the marginal effect of each variable of interest on the HRF characteristics (**Fig. 1d**), where the marginal effect is defined as the effect of a single variable on the outcome (HRF characteristics) when all other variables in the model are held at a constant value (*e.g.,* value in the original data) and finally averaged across all values and conditions, using the R package *marginaleffects*^102^.

### Model comparisons

We assessed the importance of the variables using model comparisons. A series of five candidate models were constructed, ranging from a baseline fixed effects only model to a full model containing all fixed and random effects (*type*: IED type; *sign*: HRF sign).

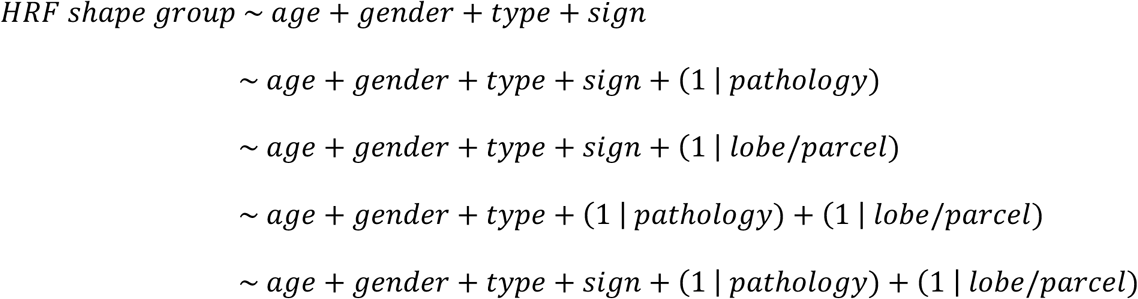

All models were fit in R using the *mclogit* package^103^, employing the penalized quasi-likelihood approximation with a maximum likelihood estimator. We selected the best fitting and most parsimonious model by ranking the models based on a comprehensive set of metrics computed via the R *performance* package^104^. These metrics included: explained variance (Nagelkerke’s pseudo-R²), model fitting error (Root Mean Squared Error, RMSE), predictive accuracy (Akaike Information Criterion, AIC), and parsimony the simplest model with the greatest explanatory power (Bayesian Information Criterion, BIC). Finally, these metrics were normalized and rescaled from 0% to 100% (where 100% represents the top performance across all models).

### Pathology classification

To further illustrate the association between pathology and HRF features, we opted to classify pathology using only six HRF features with a random forest classifier. The analysis was conducted in R, primarily using the *tidymodels* framework^105^. To obtain a robust estimate of model performance, we applied a 5×5 nested cross-validation strategy. The outer loop was used for testing the model performance, while the inner loop was used for hyperparameter tuning. Data with complex pathology type were excluded from this analysis due to insufficient representation in each cross-validation fold. To account for the imbalanced distribution of pathology classes, both inner and outer cross validation splits were stratified by the outcome variable (*i.e.,* pathology). Within each inner loop, we performed a random search over 100 combinations to tune hyperparameters. The optimal hyperparameter set was selected by maximizing the F1-score. To further address class imbalance, the training data was preprocessed by random upsampling the minority classes to 50% of the size of the majority class. The final model performance was evaluated on the hold-out sets from the outer cross validation loop. We calculated the mean and standard deviation of several metrics across the five folds, including the precision, recall, F1-score, Matthew’s Correlation Coefficient (MCC), and Cohen’s Kappa (κ). A confusion matrix was also generated by pooling the predictions from all outer folds. We also computed Gini importance to assess influence of each feature on classification performance. Finally, to estimate the null distributions of the performance metrics, we randomly permuted the pathology labels 1,000 times without replacement. The resulting distributions of each metric represent realistic chance level values.

### Robustness to choice of parcellation atlas

To assess the robustness of the results from the aforementioned analyses, we repeated them using the Schaefer2018 atlas^37^ with 200 parcel and the subcortical parcellation from FreeSurfer. Unlike the voxel-ROI-lobe-brain hierarchical structure used with the FreeSurfer segmentation, we replaced the lobe level with the network level defined by the Yeo7 networks^106^ in this analysis.

## Data and code availability

Raw data files supporting the findings of this study are available upon reasonable request and approval by the ethics boards of the corresponding institutions. Processing code is available at https://github.com/zhengchencai/epilepsy_hrf_lib. The repository includes scripts for HRF deconvolution using AFNI-based processing adapted from Chen et al., 2023^18^ available at https://github.com/afni/apaper_hrf_profiles; HRF library construction using the time decomposition method adapted from Kay et al., 2020^34^ and Allen et al., 2022^107^ available at https://github.com/cvnlab/TDM and https://github.com/cvnlab/nsddatapaper; code to reproduce result figures; and the HRF library along with the top three principal components.

## Acknowledgements

This work was supported by the Canadian Institutes of Health Research (J.G. FDN-143208) and (B.B. FDN-154298, PJT-174995, PJT-191853). Z.C is funded by FRQS Postdoctoral Fellowship, Fonds de recherche du Québec - Santé (FRQS) and MNI Jeanne Timmins Costello Fellowship. GC was supported by the NIMH Intramural Research Program (ZICMH002888) of NIH/HHS, USA. H.M.K was supported by Japan Society for the Promotion of Science (JSPS 24K12221) and Japan Agency for Medical Research and Development (AMED JP24wm0625207). We thank Dr. Kendrick Kay for releasing the code for the time decomposition method and for addressing our questions on GitHub.

## Author contributions

Z.C., J.G. and B.B. contributed to the study conception, data analysis, interpretation of results, and manuscript writing. N.V.E., T.A., H.M.K., G.C., A.K., and C.A. contributed to data analysis and interpretation of results. H.M.K. and A.K. contributed to data acquisition and EEG annotation. R.D., D.K.N., J.H., and F.D. referred patients and provided clinical information. All authors provided substantial feedback throughout the study and contributed to manuscript revision.

