## Supplementary Material for "The Hemodynamic Response Function Varies Across Anatomical Location and Pathology in the Epileptic Brain"

Zhengchen Cai, PhD

Boris Bernhardt, PhD

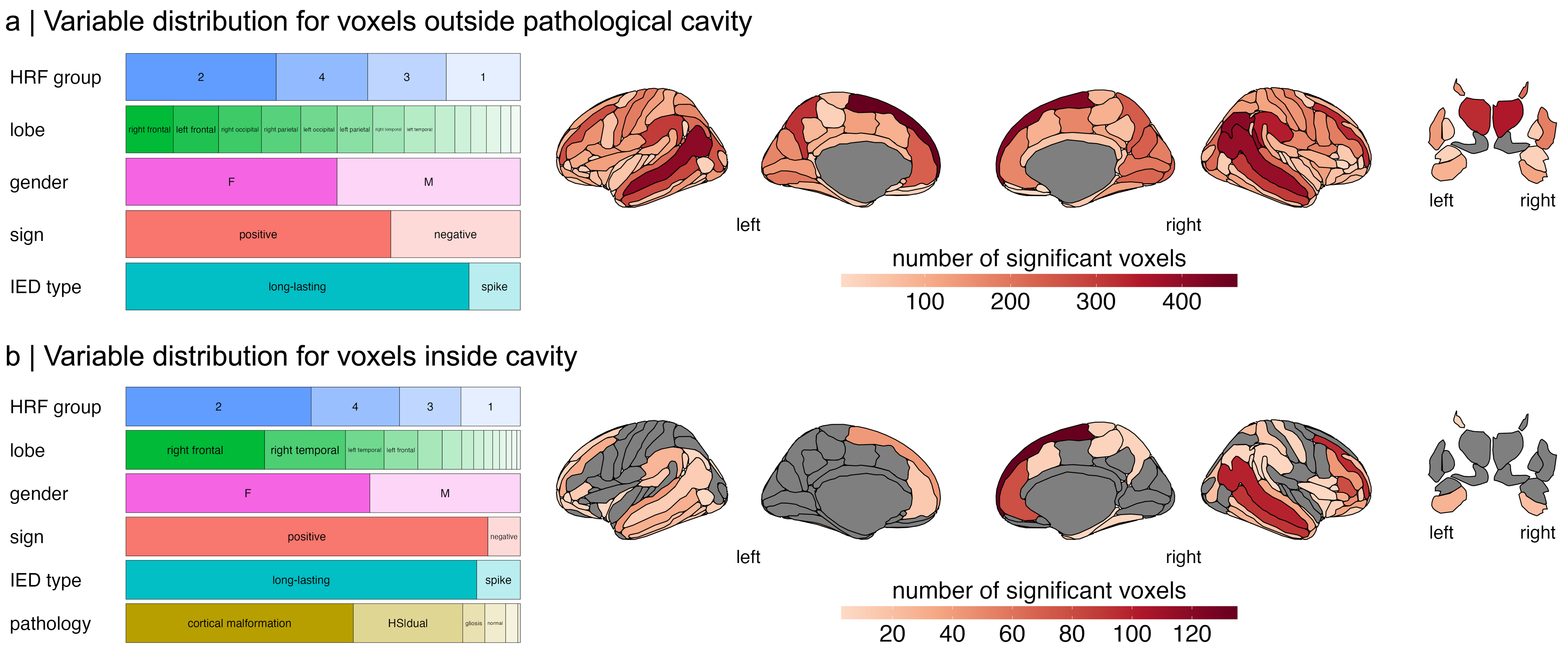

**Fig. S1 | Variable distributions in studies 2 and 3**. **a,** Variable distribution for voxels outside pathological cavity (Study 2). The distribution of voxels across lobes followed the order: frontal (23.5%), occipital (20.1%), parietal (19.0%), temporal (15.8%), subcortical (9.0%), cingulate (7.6%) and insula (5.0%). The sample was relatively balanced by gender (F: 53.5%), and by four HRF shape groups (Group 1: 18.8%, Group 2: 38.1%, Group 3: 19.9% and Group 4: 23.2%). There were more positive HRFs (67.2%) than negative ones and more long-lasting IEDs (87.0%) than spikes. **b,** Variable distribution for voxels inside surgical cavity (Study 3). The majority voxels located in the frontal (43.7%) and temporal (30.3%) lobes, followed by the parietal (7.5%), cingulate (7.1%), occipital (4.7%), subcortical (3.5%), and insular (3.2%) regions. Overall, the sample was relatively balanced by gender (F: 61.8%). There were more Group 2 HRFs (47.0%) than others (Group 1: 15.0%, Group 3: 15.6% and Group 4: 22.4%). Most HRFs were positive (91.8%) rather than negative, and the majority of IEDs were long-lasting (88.9%) rather than spikes. Pathology distribution was dominated by cortical malformation (57.7%) and HS/dual pathology (27.7%), followed by gliosis (5.6%), normal (5.3%), encephalitis (3.1%), and complex (0.6%).

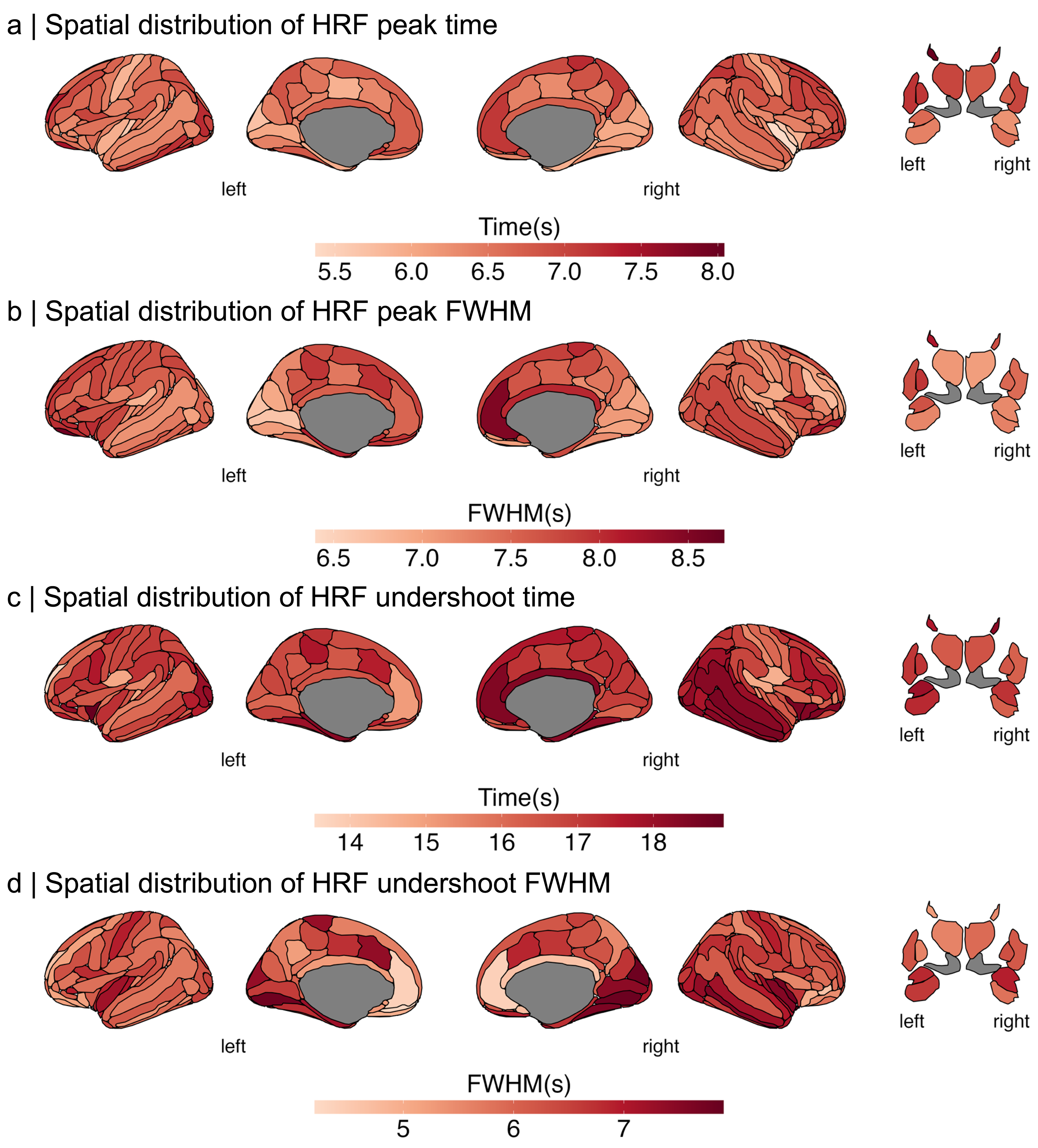

**Fig. S2 | Spatial distributions of HRF peak/undershoot times and FWHMs. a-b,** Parcel level spatial distributions of HRF peak time and full width at half maximum (FWHM). Earlier peak times (a) occurred in the primary cortices (motor, somatosensory, auditory, and visual) compared to other regions. **c-d,** Parcel level spatial distributions of HRF undershoot time and FWHM. Left-right symmetry was more consistent subcortically than cortically. Analysis included 148 cortical parcels from the Destrieux atlas and 12 subcortical ROIs from FreeSurfer segmentations.

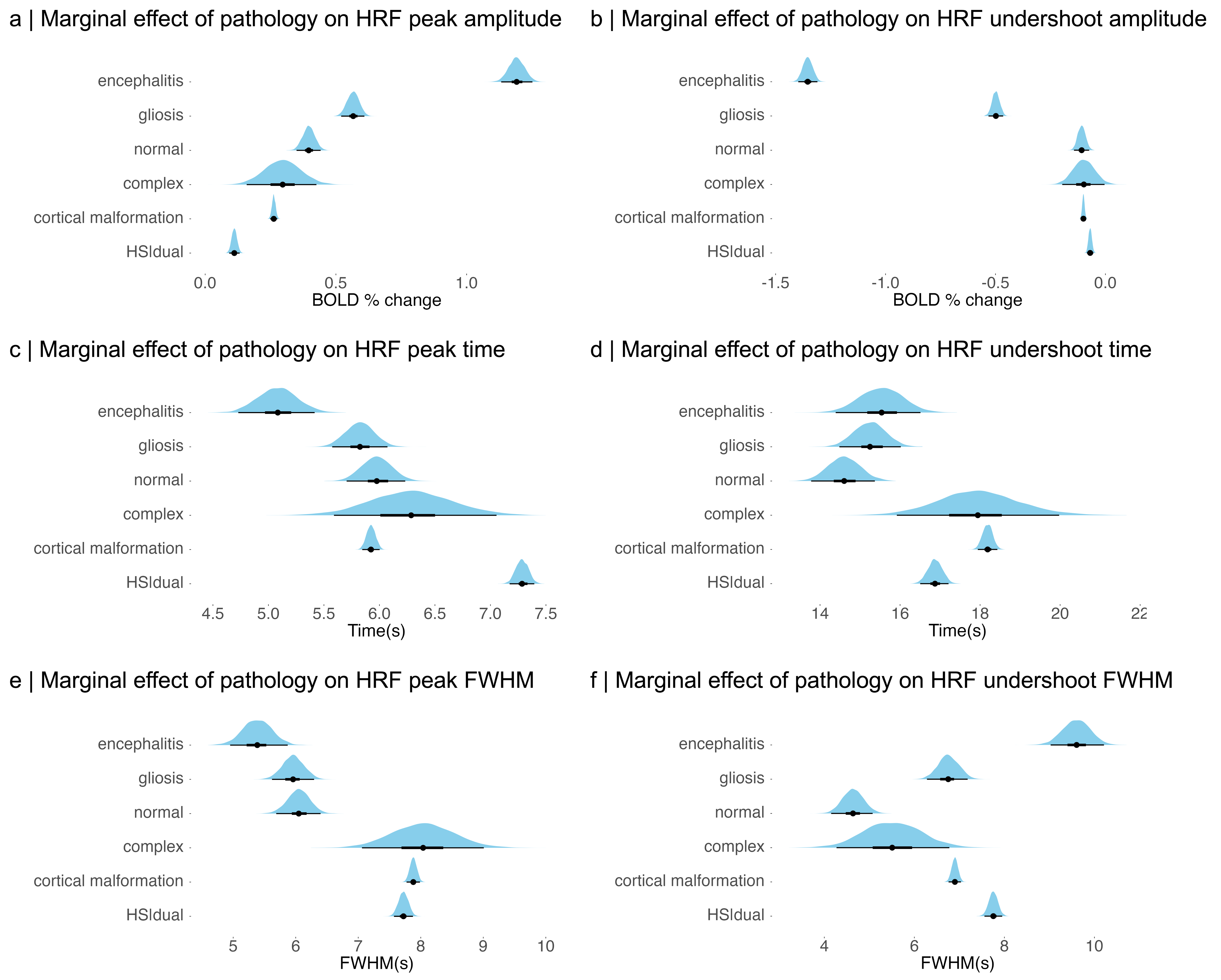

**Fig. S3 | Marginal effect of pathology on HRF undershoot features. a-b,** Marginal effect of pathology on HRF amplitudes. **c-d,** Marginal effect of pathology on HRF undershoot time to peak. **e-f,** Marginal effect of pathology on HRF undershoot full width at half maximum (FWHM). Each marginal effect is displayed as a blue density plot, with the black dot indicating the median, the thick black bar indicating the 50% highest density interval (HDI), and the thin black bar indicating the 95% HDI.

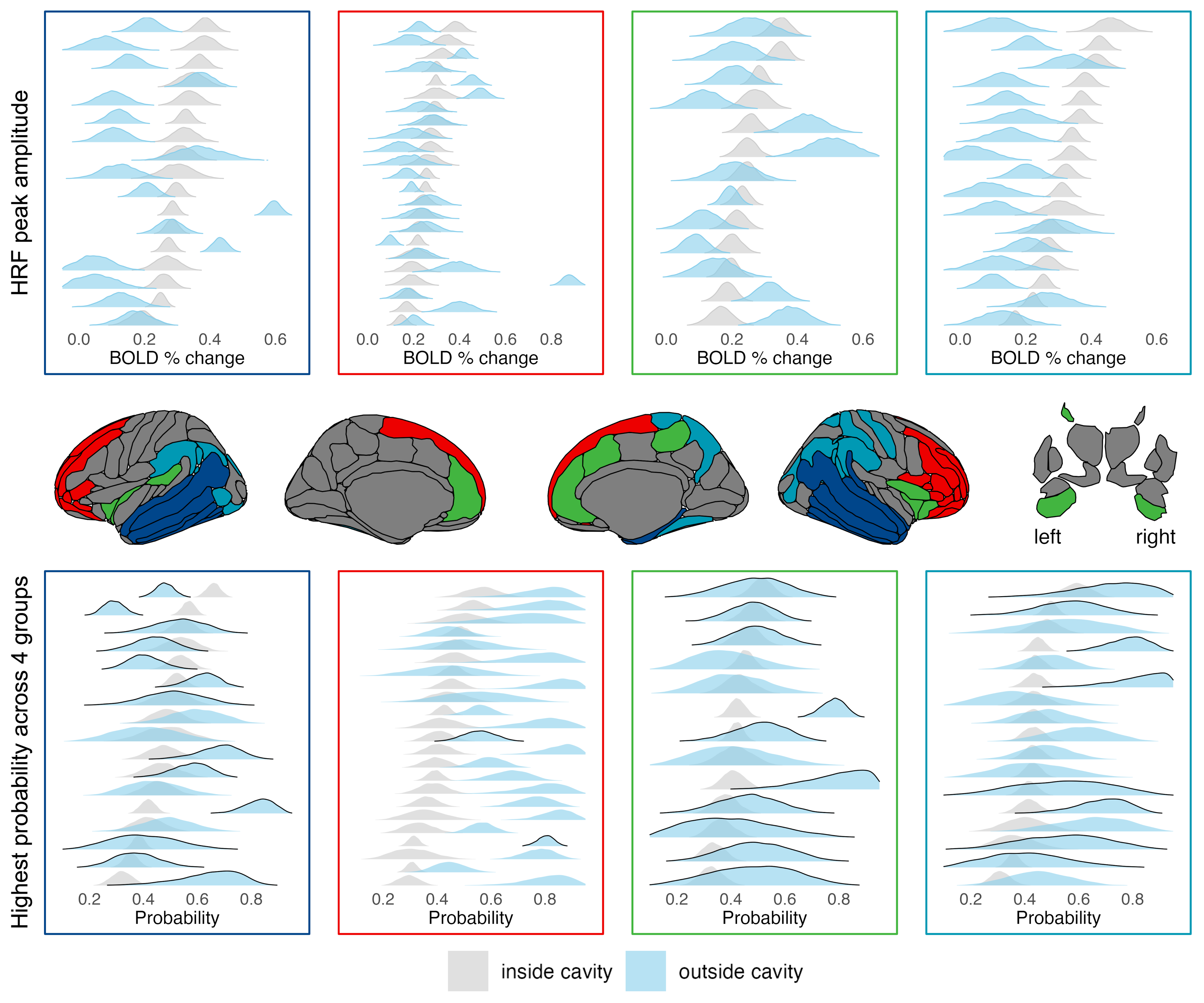

**Fig. S4 | Comparison of HRF features in the same brain regions with and without pathology.** Each blue distribution plot shows the HRF peak amplitude (top) and the highest probability across four HRF shape groups (bottom) for a given brain parcel using data with pathology (inside the surgical cavity). Parcels are grouped for visualization, with approximately equal numbers of parcels (middle panel): temporal (blue), frontal (red), insula/cingulate/subcortical (green), and parietal/occipital (cyan). The corresponding distributions from data without pathology (outside the surgical cavity) are shown in grey distributions as a normative reference. Distributions outlined in black in the bottom panel indicate parcels where the most probable HRF shape group differs between the pathological and non-pathological conditions. Clear contrasts in both HRF peak amplitude and HRF shape group are observed between with and without pathology in the same parcel.

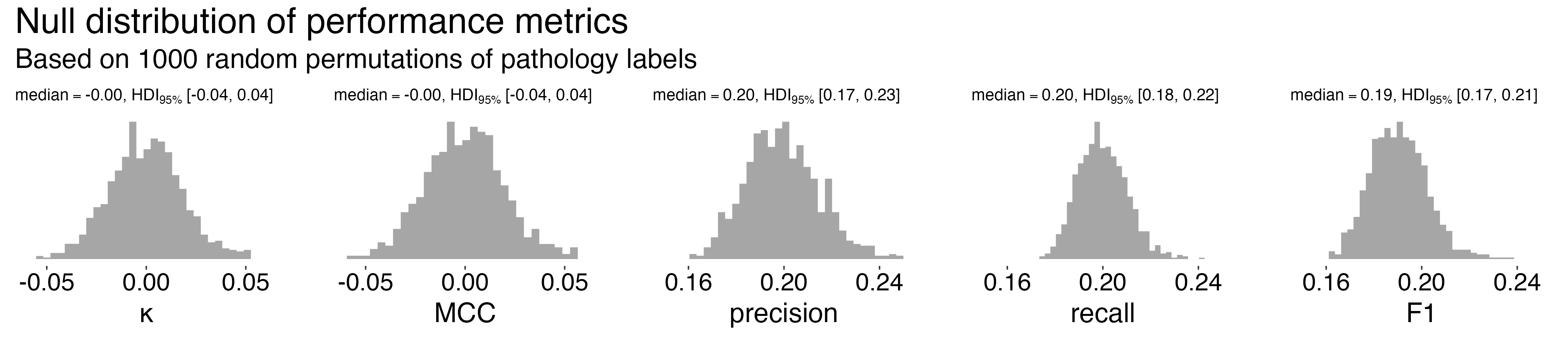

**Fig. S5 | Null distribution of pathology classification performance metrics.** Each histogram shows the null distribution of a performance metric obtained by randomly permuting the pathology labels 1,000 times (without replacement). The median and 95% highest density interval (HDI_95%_) are indicated above each histogram.

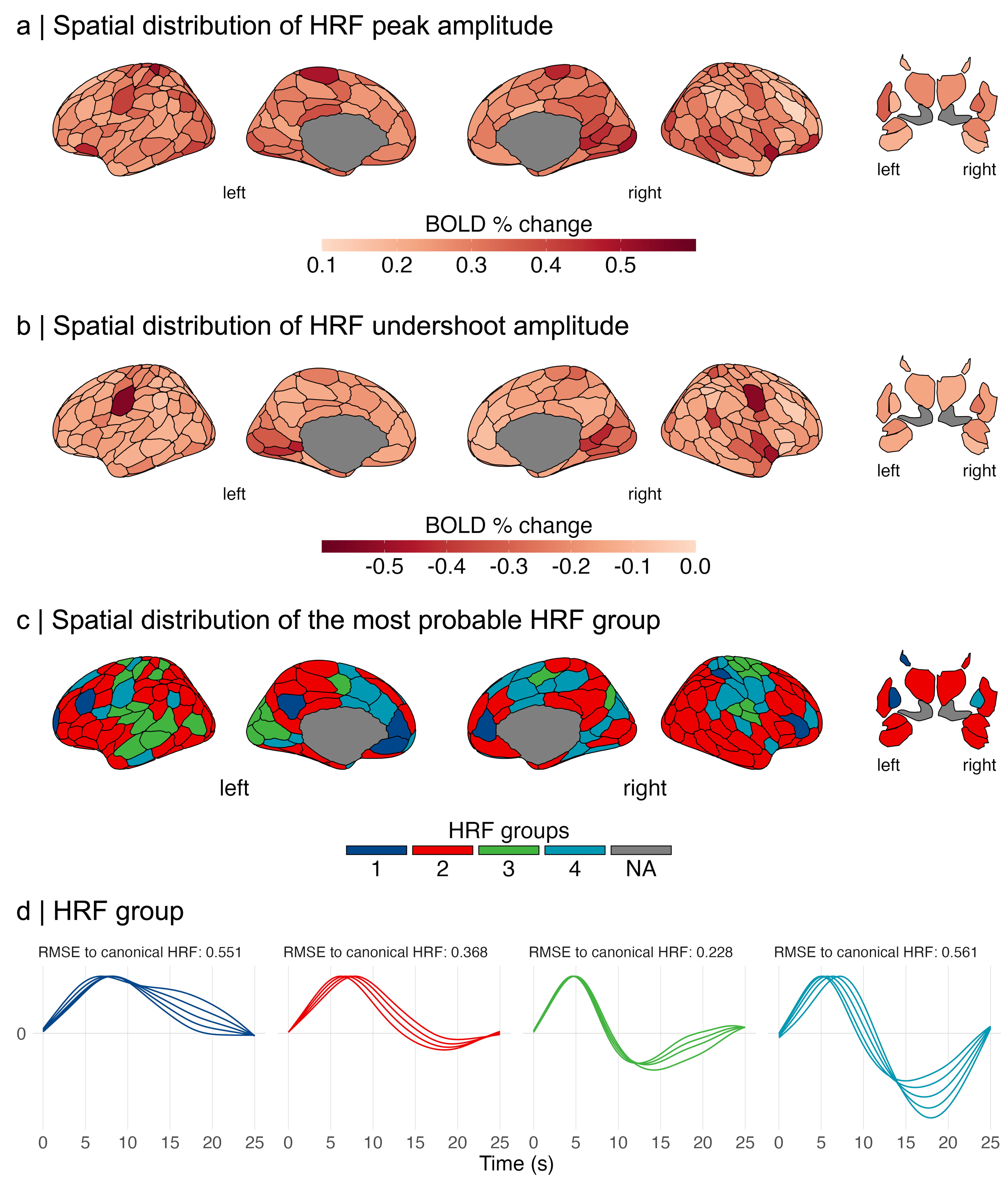

**Fig. S6 | HRF features show anatomical location association in the epileptic brain (using Schaefer2018 atlas). a-b,** Parcel level spatial distributions of HRF peak and undershoot amplitudes. Analysis included 200 cortical parcels from the Schaefer2018 atlas and 12 subcortical ROIs from FreeSurfer segmentations. Parcels within the primary cortices (motor, somatosensory, auditory, and visual) exhibited larger amplitude magnitudes compared to other regions. **c,** Parcel level spatial distribution of the most probable HRF shape group (i.e., the group with the highest marginal effect for each parcel). The same primary vs. others cortex pattern observed in (a) is also apparent here. **d,** HRFs in the library were classified into four shape-based groups: Group 1 (no undershoot), Group 2 (low undershoot), Group 3 (moderate and early undershoot), and Group 4 (large and late undershoot). The average root mean square error (RMSE) relative to the canonical HRF was 0.551, 0.368, 0.228, and 0.561, respectively.

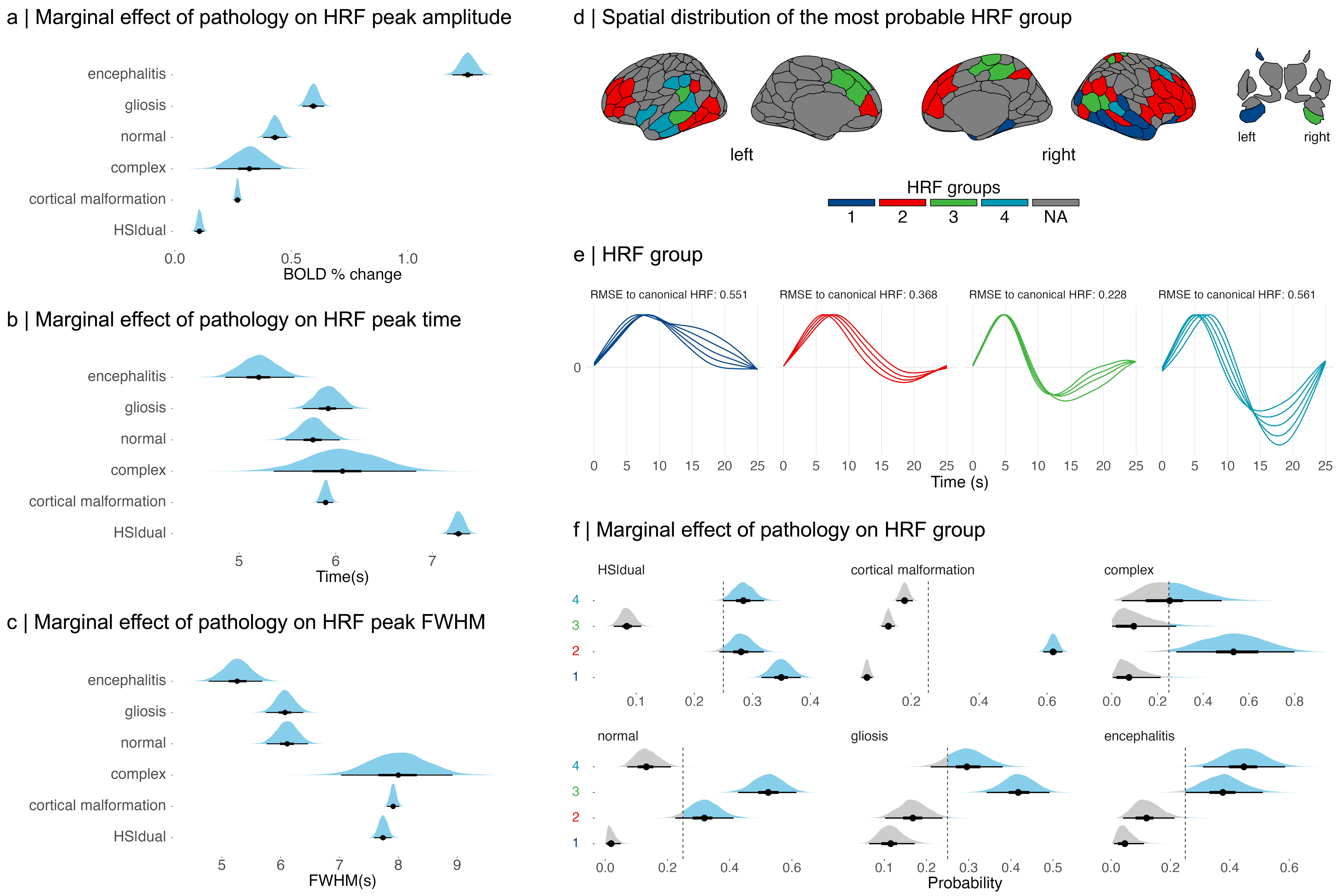

**Fig. S7 | HRF features show association with pathology in epileptic brain (using Schaefer2018 atlas). a-c,** Parcel level spatial distributions of HRF peak amplitude, time to peak, and full width at half maximum (FWHM). The normal pathology type exhibited mid-range values, separating values observed in gliosis and encephalitis from those in HS/dual (*i.e.,* hippocampal sclerosis plus another pathology) and cortical malformation. **d,** Parcel level spatial distribution of the most probable HRF shape group (i.e., the group with the highest marginal effect per parcel). HRF shape groups showed pathology specific spatial distributions. **e,** The HRF library and corresponding HRF shape groups. **f,** Marginal effects of pathology on HRF shape group. Vertical dashed lines indicate the 25% chance probability across the four groups. Areas below this reference are shaded in gray. Each marginal effect is displayed as a blue density plot, with the black dot indicating the median, the thick black bar indicating the 50% highest density interval (HDI), and the thin black bar indicating the 95% HDI.

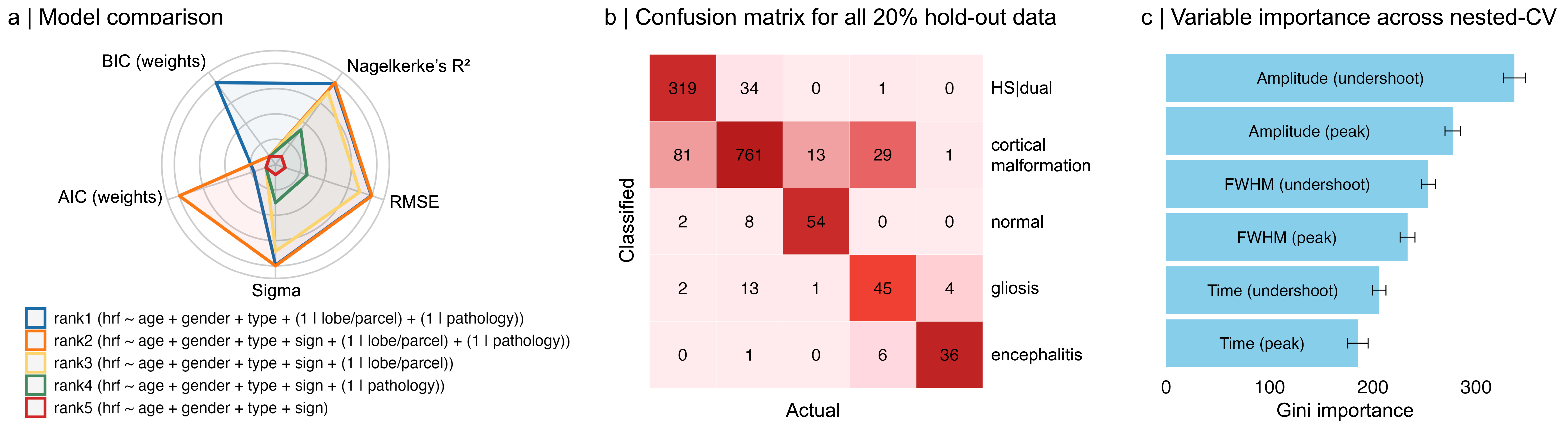

**Fig. S8 | Associations between HRF features and pathology revealed by model comparison and classification (using Schaefer2018 atlas). a,** A radar plot shows model comparison results across five candidate models, ranging from a baseline fixed effects only model to a full model including all fixed and random effects. Each axis represents one model performance metric: Nagelkerke's pseudo-R², root mean squared error (RMSE), Akaike information criterion (AIC), and Bayesian information criterion (BIC). Each metric was normalized and rescaled from 0% to 100% relative to the best performance across models. Models were ranked by overall performance across all metrics. **b,** The confusion matrix from pathology classification using 20% hold-out data. The highest values appear along the diagonal, indicating accurate classification. **c,** Gini importance of HRF features from the classification model. Feature importance decreased from amplitude to FWHM and then to time, with undershoot related measures showing slightly higher values within each category. Black error bars indicate the standard deviation across cross validation folds.

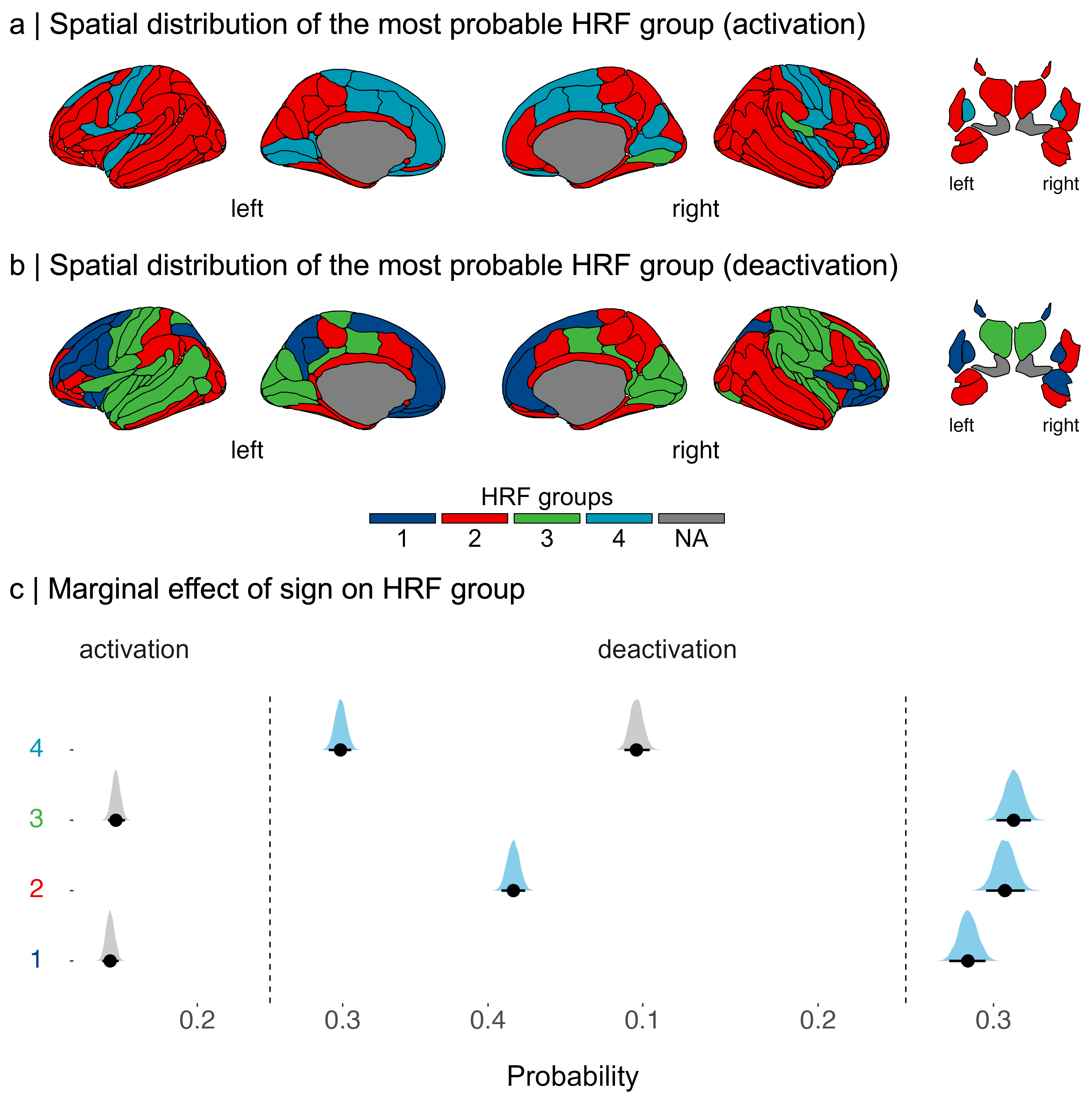

**Fig. S9 | HRF features show anatomical location association in epileptic brain when stratified by HRF sign. a,** Parcel level spatial distribution of the most probable HRF shape group (*i.e.,* the group with the highest marginal effect for each parcel) for positive (activation) HRFs. **b,** Parcel level spatial distribution of the most probable HRF shape group for negative (deactivation) HRFs. The primary vs. others cortex pattern was apparent in both (a) and (b). Analysis included 148 cortical parcels from the Destrieux atlas and 12 subcortical ROIs from FreeSurfer segmentations. **c,** Marginal effect of HRF sign on HRF group. Positive HRFs were primarily in Group 2 and Group 4, with the remaining voxels evenly divided between Groups 1 and 3. Negative HRFs were similarly distributed across Groups 1-3, with a low probability of being Group 4. Vertical dashed lines indicate the 25% chance probability across the four groups. Areas below this reference are shaded in gray. Each marginal effect is displayed as a blue density plot, with the black dot indicating the median, the thick black bar indicating the 50% highest density interval (HDI), and the thin black bar indicating the 95% HDI.

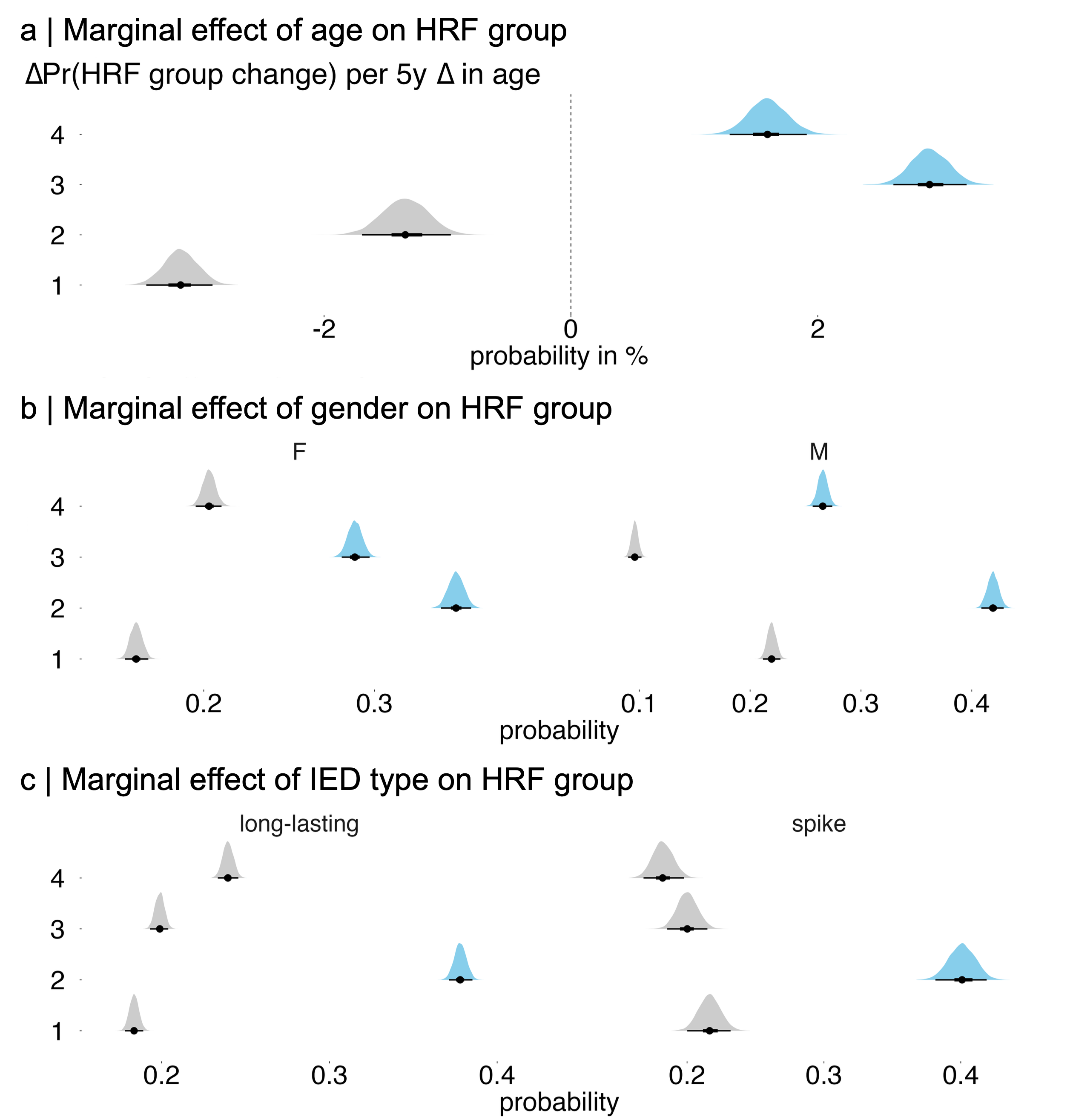

**Fig. S10 | Effects of other factors on HRF are subtle. a,** Marginal effect of age on HRF group. The absolute probability change of belonging to a given HRF group with a 5-year increase in age was always less than 4%, with the largest effect observed in Group 1. Vertical dashed lines indicate the 0% probability change for any HRF group. Areas below this reference are shaded in gray. **b,** Marginal effect of gender on HRF group. Both females and males showed the highest probability for HRF Group 2. The difference was mainly in the secondary groups where males showed a higher probability for Group 4. **c,** Marginal effect of IED type on HRF group. Both long-lasting IEDs and spikes were most frequently associated with Group 2, with the other three groups each accounting for approximately 20%. Areas below 25% chance reference are shaded in gray. Each marginal effect is displayed as a blue density plot, with the black dot indicating the median, the thick black bar indicating the 50% highest density interval (HDI), and the thin black bar indicating the 95% HDI.
